# Leveraging probability concepts for genotype by environment recommendation

**DOI:** 10.1101/2021.04.21.440774

**Authors:** Kaio O.G. Dias, Jhonathan P.R. dos Santos, Matheus D. Krause, Hans-Peter Piepho, Lauro J.M. Guimarães, Maria M. Pastina, Antonio A.F. Garcia

## Abstract

Statistical models that capture the phenotypic plasticity of a genotype across environments are crucial in plant breeding programs to potentially identify parents, generate offspring, and obtain highly productive genotypes for distinct environments. In this study, our aim is to leverage concepts of Bayesian models and probability methods of stability analysis to untangle genotype-by-environment interaction (GEI). The proposed method employs the posterior distribution obtained with the No-U-Turn sampler algorithm to get Monte Carlo estimates of adaptation and stability probabilities. We applied the proposed models in two empirical tropical datasets. Our findings provide a basis to enhance our ability to consider the uncertainty of cultivar recommendation for global or specific adaptation. We further demonstrate that probability methods of stability analysis in a Bayesian framework are a powerful tool for unraveling GEI given a defined intensity of selection that results in a more informed decision-making process towards cultivar recommendation in multi-environment trials.

## Introduction

In plant breeding, the evaluation of crops in multi-environmental trials (MET) is a critical step to understand their stability and overall performance for data-driven decisions. The different responses of genotypes across distinct environments triggered by genotype-by-environmental interaction (GEI) are a well-known challenge in crop improvement (Crossa 1990; Falconer and Mackay 1996). Understanding the genetic basis of GEI has been an open field for many decades. A century ago, RA Fisher described the GEI as a non-linear interaction (biometric concept), while Lancelot Hogben described it as a third class of variability (developmental concept), which arises from genetic and environmental compositions (Tabery, 2008). From the Fisher-Hogben debate to the present, several statistical models to deal with GEI under a variety of assumptions have been proposed (Yates and Cochran, 1938; Crossa, 1990; Smith et al., 2005; Crossa, 2012; Malosetti et al., 2013, Mackay et al., 2019).

GEI can be either ignored or explored over the selection process. When recommendations ignore GEI, the crop adaptation can potentially be lost and might compromise the yield potential in specific environments (Malosetti et al. 2013; Yan 2016; Gage et al. 2017). The exploration of GEI allows for the maximum expression of the yield potential, triggered by adaptive responses of cultivars to a target population of environments. By exploring GEI, stability analysis can provide information on the norms of reaction of a genotype in different environments. Early models estimate the stability of genotypes by the regression of their means on an environmental index (e.g., the site means) (Finlay and Wilkinson 1963; Eberhart and Russell 1966), or by considering the stability variance (Shukla 1972). Later on, multivariate models such as principal components analysis (PCA) of residuals from two-way table ANOVA (usually known as an additive main effect and multiplicative interaction (AMMI), Gauch and Zobel 1988), PCA of genotype plus genotype-by-environment analysis (so-called GGEBiplot, Yan et al. 2000), site regression model (Crossa and Cornelius 1997), and factor analytic multiplicative mixed model (Piepho 1997; Smith et al. 2001) have become widely used for stability analysis and cultivar recommendation. The extension of these models has also been proposed in a Bayesian framework (Cotes et al. 2006; Crossa et al. 2011; Josse et al. 2014; Da Silva et al. 2015).

The importance of developing statistical models that explore GEI and are informative about genotype stability can potentially improve crop adaptation. The phenotypic plasticity of a genotype across environments is therefore crucial to identify parents for future breeding generations, generate superior offspring, and obtain highly productive genotypes for distinct environments. Currently, linear mixed models and hierarchical Bayesian models are commonly used to deal with GEI. These models are highly desirable due to their capacity to take into account complex plant breeding scenarios with unbalanced data, heterogeneous covariances, and spatial field trends (Smith et al. 2005; Crossa 2012; Malosetti et al. 2013). Furthermore, these models allow incorporating relationships among individuals (Piepho et al. 2008; Burgueño et al. 2012; Gage et al. 2017; Dias et al. 2018) and environmental variables for a reaction norm model (van Eeuwijk et al. 1996; Li et al. 2018; Millet et al. 2019; Costa-Neto et al. 2020). These advantages enable a better understanding of the GEI’s underlying mechanisms, and to predict new genotypes in yet-to-be-seen environments.

Despite the importance of the outlined methodologies, interpreting their results might not be trivial with substantial amounts of genotypes, trials, and cycles of selection frequently observed in a breeding program. Most studies of GEI rely on graphical representations out of results from regression analysis. Although these procedures have a prominent role in crop improvement to identify GEI patterns, with popular methodologies such as biplot analysis (Gabriel 1971), it is still a challenge to visualize and derive inference from trials with many genotypes, locations, and years. Furthermore, the projection of GEI in the Cartesian plane can also not fully reflect the complex variation in multidimensional data space. Novel methodologies with unified, straightforward visualization, and measures of uncertainty of phenotypic response to environmental gradients are critical to better improve crop adaptation and cultivar recommendation (Smith and Cullis 2018).

As mentioned by Wallace et al. (2018), a central question that needs to be answered to advance breeding is: “How do we adapt crops to better fit agricultural environments?”. The application of probability-base methods could be a crucial stepping stone to enhance our ability to identify stable cultivars across a wide variety of distinct environments or breeding regions. The Bayesian framework is particularly interesting to address this question because of its capacity to integrate knowledge from previous experiments into priors (Crossa et al. 2011; Gelman et al. 2013; dos Santos et al. 2020) and by exploring GEI (Crossa et al. 2011; Josse et al. 2014; de Oliveira et al. 2015) and stability variances (Edwards and Jannink 2006). The goal of our study is therefore to propose analytical strategies based on Bayesian models that take full advantage of posterior probabilities to facilitate the decision-making process towards cultivar recommendation in a commercial breeding program. To the best of our knowledge, no previous study has explored these probability methods in studying GEI and in cultivar recommendations, and usually, most of them restrict inference only to biplot analysis. Our approach can be applied for both global and specific cultivar recommendation/adaptation in the target population of environments or breeding regions. Its main advantage is that probabilistic statements are conveniently derived from the posterior samples that combine mean performance and stability. We also present step-by-step theoretical concepts to easily implement the proposed strategies in a MET dataset. The proposed approach is illustrated in two empirical datasets of maize (*Zea mays* L.) and wheat (*Triticum aestivum* L.). The developed R code and the dataset to reproduce all the analyses are publicly available.

## Materials and Methods

### Motivational and Theory

To introduce terminology and definitions of the proposed models, consider that we have the marginal genetic value of the overall performance of *j*^*th*^ genotype in *k*^*th*^ environments for a given breeding zone (Table 1). Each cell represents the realization of the *s*^*th*^ sample from the posterior distribution (*p*(*g*|*y*)) of the genetic value of the *j*^*th*^ genotype 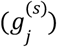 given its observed phenotype *y* (Table 1).

**Table 1.**
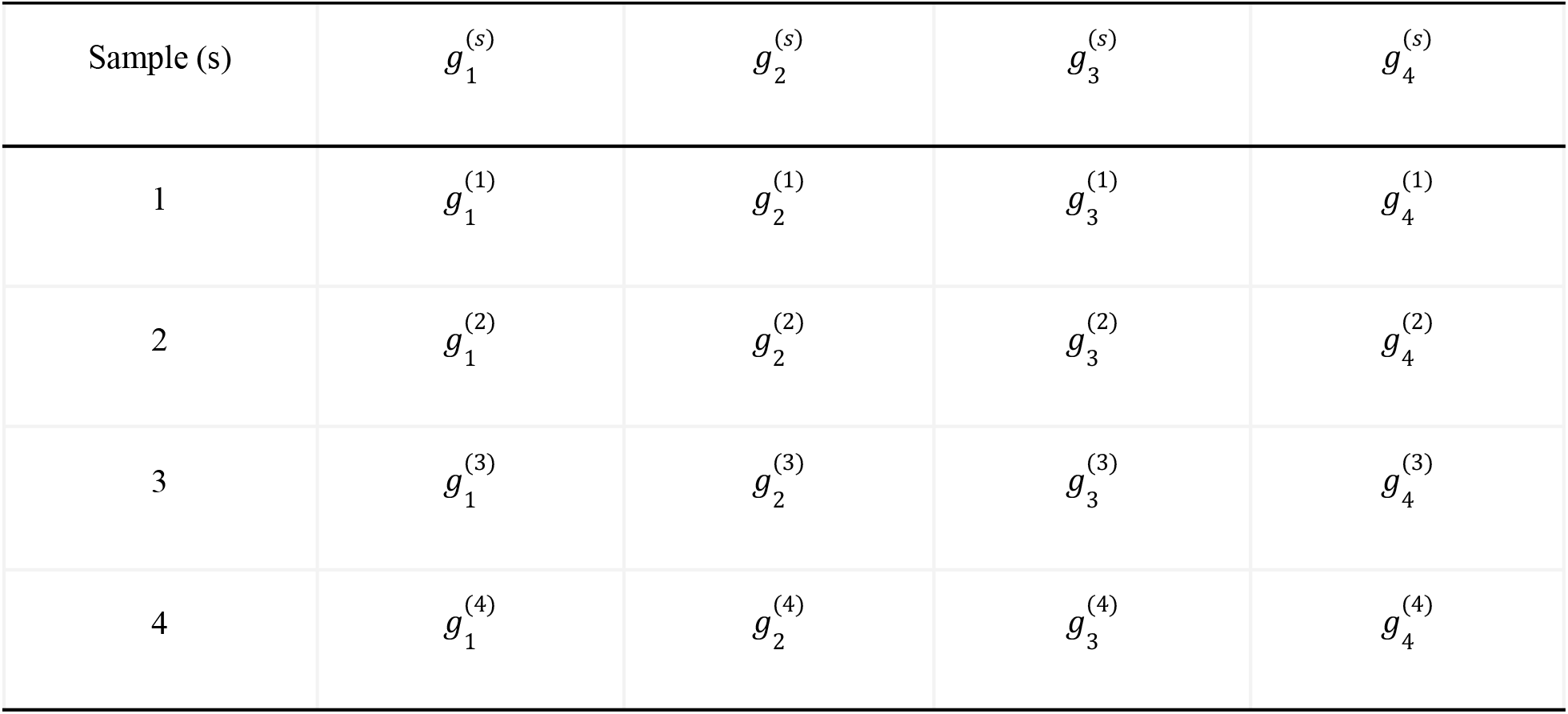
Hypothetical scheme of the marginal performance of *j*^*th*^ genotype 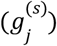 in *k*^*th*^ environments from the *s*^*th*^ sample from the posterior distribution.

| Sample (s) | $g_1^{(s)}$ | $g_2^{(s)}$ | $g_3^{(s)}$ | $g_4^{(s)}$ |
| --- | --- | --- | --- | --- |
| 1 | $g_1^{(1)}$ | $g_2^{(1)}$ | $g_3^{(1)}$ | $g_4^{(1)}$ |
| 2 | $g_1^{(2)}$ | $g_2^{(2)}$ | $g_3^{(2)}$ | $g_4^{(2)}$ |
| 3 | $g_1^{(3)}$ | $g_2^{(3)}$ | $g_3^{(3)}$ | $g_4^{(3)}$ |
| 4 | $g_1^{(4)}$ | $g_2^{(4)}$ | $g_3^{(4)}$ | $g_4^{(4)}$ |

Suppose now that we can rank the genotypes in each row of Table 1 and create indicator variables (*I*) representing the success of a given genotype belonging to a subset of superior genotypes (Ω) under a predefined selection intensity (e.g., the 20% higher performance genotypes), given by 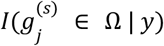, by considering the posterior samples (Table 2).

**Table 2.** Hypothetical scheme of indicators mapping if the marginal genetic value of *j*^*th*^ genotype belongs (1) or not (0) to a subset of superior genotypes (Ω) predefined by the breeder, that is, 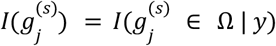.

| Sample (s) | $I(g_1^{(s)})$ | $I(g_2^{(s)})$ | $I(g_3^{(s)})$ | $I(g_4^{(s)})$ |
| --- | --- | --- | --- | --- |
| 1 | 1 | 1 | 1 | 0 |
| 2 | 1 | 1 | 0 | 0 |
| 3 | 1 | 0 | 1 | 0 |
| 4 | 0 | 0 | 0 | 1 |
| $\sum_{s=1}^S I(g_j^{(s)})$ | 3 | 2 | 2 | 1 |

We can compute the probability with the discretized samples (e.g. Monte Carlo sampling) mentioned in Table 1 by the ratio of success (being or not in Ω) and the total number of events (total number of posterior samples) (Gelman et al. 2013). Thus, the probability of a genotype belonging to a subset of superior genotypes (probability of performance) can be calculated by:

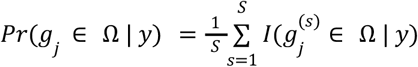

where 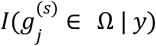 is an indicator variable mapping success (1) if the 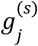 sample is in the subset Ω and failure (0) otherwise, and S the total number of samples.

To illustrate a hypothetical example displaying how we can use the structure of Table 2 to compute the probability of performance (*Pr*(*g*_*j*_ ∈ Ω | *y*)) above, consider genotype 1. This probability would be 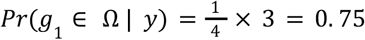, which means there is a probability of 75% of the marginal genetic value of genotype 1 being among the superior genotypes evaluated in *k*^*th*^ environments, for a given breeding zone, grounded on the predefined selection intensity. A given breeding zone is represented by a sample of trials in that geography.

The pairwise relationship of the marginal probability of performance among genotypes can also give a global comparative picture for selection to a wide variety of environments. We might be able to understand such a relationship by computing the probability that the *i*^*th*^ genotype has a better performance than the *j*^*th*^ genotype, when evaluated in *k*^*th*^ environments, by considering their estimated genetic values *g*_*i*_ and *g*_*j*_, respectively. The pairwise probability of performance can be obtained by,

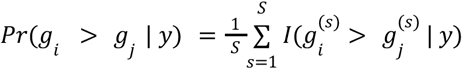

where 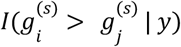 is an indicator variable mapping success (1) if the posterior sample 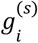 is greater than 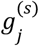 and failure (0) otherwise. Our marginal pairwise probability has an analogy to the Plackett-Luce model (Plackett 1975), which relies on Luce’s axiom of choice (Luce 1977), for modeling ranking data. As stated by van Etten et al. (2019) the axiom states that the odds of one item beating another does not depend on any other items from which the choice was made.

Suppose now we want to compute the probability of performance for a specific environment or breeding zone. Table 3 shows a hypothetical scheme illustrating how we can arrange in a three-dimensional array the indicators 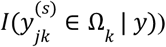 mapping 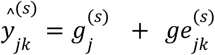, the expected performance of *j*^*th*^ genotype belongs to the superior set of genotypes in environment or breeding zone k (Ω_*k*_).

**Table 3.** Hypothetical scheme of the expected predicted performance 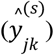 of the genotype *j* evaluated on *k* environments (Env) from the *s*^*th*^ sample from the posterior distribution, where 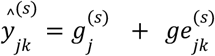.

|  | Env 1 |  | Env 2 |  | Env 3 |  |
| --- | --- | --- | --- | --- | --- | --- |
| Sample (s) | $I(y_{11}^{(s)})$ | $I(y_{21}^{(s)})$ | $I(y_{12}^{(s)})$ | $I(y_{22}^{(s)})$ | $I(y_{13}^{(s)})$ | $I(y_{23}^{(s)})$ |
| 1 | 1 | 1 | 0 | 0 | 1 | 0 |
| 2 | 0 | 1 | 1 | 0 | 1 | 1 |
| 3 | 0 | 1 | 0 | 1 | 0 | 1 |
| 4 | 0 | 0 | 1 | 0 | 0 | 1 |
| $\sum_{s=1}^S I(y_{jk}^{(s)})$ | 1 | 3 | 2 | 1 | 2 | 3 |

In the latter case, we can estimate the conditional probability of successfully selecting a given genotype to a particular environment or breeding zone (the probability of performance within the environment) by,

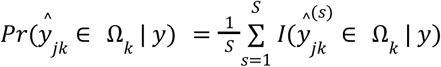

For the case of the genotype 1 evaluated at environment 3 for example, this probability would be given by 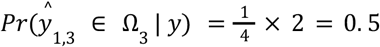. This means that there is a probability of 50% of genotype 1 being among the superior genotypes at environment 3 based on the predefined selection intensity. On the other hand, for genotype 2, 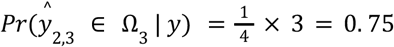.

The probability of stability of a genotype-specific across environments can be computed by the variance of GEI effects 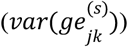. We can compute the probability of a *j*^*th*^ genotype evaluated in the *k*^*th*^ environment belonging or not to a group of genotypes with lower 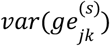, given the predefined selection intensity (Ω), by

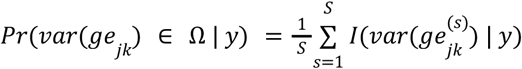

Genotypes with a lower variance of GEI tend to be more stable, i.e. the performance of a specific genotype is more predictable across environments. This measure of GEI variance for a given genotype has an analogy with the method proposed by Shukla (1972). The main difference is that our approach does not take into account the error variance (homogeneous or heterogeneous), which in Shukla (1972) is confounded with *var*(*ge*_*jk*_).

Assuming independence between genotype main effect and variance of GEI, we can compute the joint probability of performance and stability by multiplying the marginal probability of performance (*Pr*(*g*_*j*_ ∈ Ω | *y*)) and the probability of stability (i.e. genotype-specific variance) *Pr*(*var*(*ge*_*jk*_) ∈ Ω | *y*), that is,

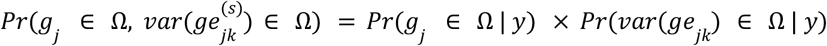

With the outlined probability statements, we further addressed a major concern faced by farmers and plant breeders to minimize the risk of a cultivar’s failure (Mead et al. 1986; Piepho 1996), implicating financial loss for farmers and a possible reduction in market share. To quantify the risk in cultivar recommendation, Eskridge (1990) presented a safety-first index. This index has a connection with our proposal, once it combines information about mean performance and stability variance. The great advantage of Bayesian models is that, once we have fitted the model, the posterior distribution encoded all the information about the predicted parameters. An extension of the safety-first index was therefore proposed using the posterior information from the fitted model to quantify the uncertainty in cultivar recommendation.

### Empirical maize hybrid dataset

The maize dataset comprised 36 single-cross hybrids from the late-stage breeding of Embrapa Maize in Sorghum. These hybrids were evaluated for grain yield in 16 environments across five tropical regions (MN, NE, TA, TA, and TR) in Brazil in 2013 (Table S1).

Among these hybrids, 32 were from the previous step of their breeding program plus four commercial checks. After these late-stage trials, normally 8 hybrids are selected for testing the value for cultivation and use (VCU). This will be the intensity of selection (25%).

In each environment, trials were laid out as incomplete block design, using a block size of 6 and two replications per trial. The plot size was 8 square meters. Agronomic practices were performed as recommended for maize in each region of Brazil.

### Fitting a Bayesian model for the maize dataset

Four different Bayesian models were proposed in our study for partitioning the GEI variation. The first model (henceforth called M1) has a conditional normal likelihood as follows:

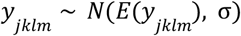

where:

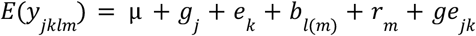

where *y_jklm_* is the phenotypic value of the *j^th^* (*j* = 1, 2,…, 36) genotype evaluated in the *l^th^* (*l* = 1, 2,…, 6) block nested within *m^th^* (*m* = 1, 2) replication and *e^th^* (*k* = 1, 2,…, 16) environment; μ is the overall mean, *g_j_* the genetic effect, *b_l(m)_* the block within replication effect, *r*_*m*_ the replication effect and *ge*_*jk*_ the genotype-by-environment interaction effect.

The prior probability distribution for each parameter for the M1 model can be defined as:

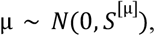

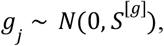

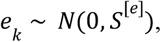

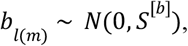

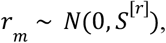

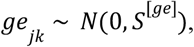

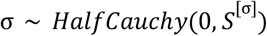

where *N*(0, *S* ^[θ]^) and *HalfCauchy*(0, *S*^[θ]^) represent a normal and half-Cauchy distributions centered at zero with different *S* ^[θ]^ scale hyperparameters, respectively. The half-Cauchy distribution is constrained to be positive. To eliminate the subjective in the choice of the hyperparameters, we also consider hyperpriors as follows,

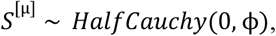

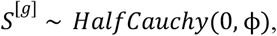

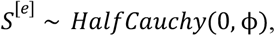

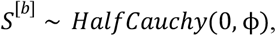

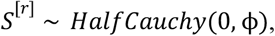

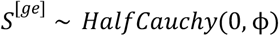

Where ϕ is the known global hyperparameter defined as ϕ = *max*(*y*)×10, defined in such a way that results in weakly informative second-level hyperpriors that allows the data to dominate the posterior distribution if the likelihood is strong (Gelman et al. 2013).

Model 2 (henceforth called M2) differs from M1 by considering a heterogeneous standard deviation of the likelihood for each environment. To this end, the prior used was: 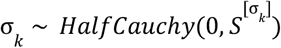. Model 3 (M3) and Model 4 (M4) differs from M1 and M2 by accounting for the effect of breeding regions (*z*_*t*_). M4 differs from M3 by considering a heterogeneous standard deviation of the likelihood for each environment while M3 considers a homogeneous standard deviation. The additional priors for M3 and M4 for the effect of breeding regions can be defined as follows:

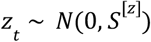

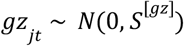

where *z* is the region effect and *gz* is the genotype-by-region interaction. All other terms were as previously described in M1.

### Bayesian safety-first index

We estimated the safety-first index proposed by Eskridge (1990) for a fixed model as:

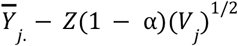

where 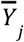 is the adjusted mean for genotype *j*across environments, *V*_*j*_ is a measure of stability (i.e. variance of GEI across environments for a given genotype) and *Z*(1 − α) is the percentile from the standard normal distribution for a value of α.

The extension of the safety-first index (Eskridge 1990) in a Bayesian framework was computed by considering the maximum a posteriori estimation of M4 for genotype main effect (*g*_*j*_), genotype-by-environment interaction (*ge*_*jk*_), and genotype-by-region interaction (*gz*_*jt*_). Also, we used the posterior distribution of the genotype main effect to estimate the percentile *Z*_*e*_(1 − α) threshold for α= 0.05. Thus, the Bayesian Eskridge’s Risk index was estimated as:

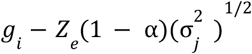

where 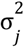 is the GEI variance (*ge*_*jk*_ + *gz*_*jt*_) for genotype *j*. All other terms were previously described.

### Empirical wheat dataset

This data set is publicly available in Crossa et al. (2010). In summary, the wheat data comprised 599 lines developed by CIMMYT evaluated in four environments for grain yield. Breeding regions were not available in this data set. Adjusted means standardized to a unit variance within each environment were made available. Lines were genotyped using Diversity Array Technology, being 1279 molecular markers after quality control.

### Fitting a Bayesian model for the wheat dataset

To evaluate our proposal considering molecular markers based on probability concepts, we assumed the four environments as independent and fitted a Bayesian ridge regression model (hereafter called M5) and computed genomic estimated breeding values (GEBV), as follows:

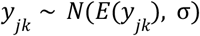

and:

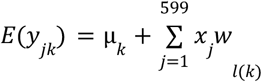

where *y*_*jk*_ is the GEBV of the *j*^*th*^ (*j* = 1, 2,…, 599) line evaluated in the *k*^*th*^ (*l* = 1, 2, 3, 4) environment; μ_*k*_ is the environmental-specific intercept, *x*_*j*_ is a column vector of marker genotypes, and *w*_*l*(*k*)_ is the estimated effect of the *l^th^* (*l* = 1, 2,…, 1279) marker in the *k*^*th*^ environment.

The prior probability distribution for each parameter for the M5 model can be defined as:

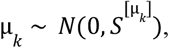

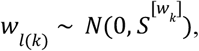

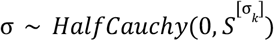

As described above, the 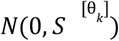 and 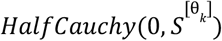 represent normal and half-Cauchy distributions centered at zero with 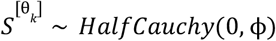, being the different 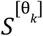 scale hyperparameters, and ϕ the known global hyperparameter. The posterior probabilities distributions were also obtained using the probabilistic programming language Stan as previously described.

### Model selection

A general description of the different models can be seen in Table 4. The posterior probability distributions of the model parameters were obtained using the probabilistic programming language Stan (Hoffman and Gelman 2014; Carpenter et al. 2017) in the interface RStan (R Core Team, 2020). For each model, four chains with 2,000 samples were fitted and 50% of the initial samples discarded as burn-in, resulting in a total of 4000 samples used for downstream analysis to compute the proposed posterior probabilities. The potential scale reduction factor 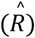, which measures the between and within chain estimates for model parameters, was used to assess the quality of the Markov chain Monte Carlo (MCMC) samples. If the 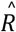 is close to one, it indicates better convergence of the samples from different chains in the parameter space. On the other hand, if chains do not mix well, 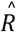 is larger than 1 (Gelman et al. 2013, pg 285).

**Table 4.** Summary of the four Bayesian models fitted in the maize data set.

| Model | Standard deviation | Expectation |
| --- | --- | --- |
| M1 | Homogeneous ($S^{[\sigma]}$) | $E(y_{jkli}) = \mu + g_j + e_k + b_{l(i)} + r_i + ge_{jk}$ |
| M2 | Heterogeneous ($S^{[\sigma_k]}$) | $E(y_{jkli}) = \mu + g_j + e_k + b_{l(i)} + r_i + ge_{jk}$ |
| M3 | Homogeneous ($S^{[\sigma_{gz}]}$) | $E(y_{jklit}) = \mu + g_j + e_k + b_{l(i)} + r_i + ge_{jk} + z_t + gz_{jt}$ |
| M4 | Heterogeneous ($S^{[\sigma_{gz_k}]}$) | $E(y_{jklit}) = \mu + g_j + e_k + b_{l(i)} + r_i + ge_{jk} + z_t + gz_{jt}$ |

$\mu$ represents the overall mean; $g_j$ genotypes; $e_k$ environments; $b_{l(i)}$ blocks; $r_i$ replications; $ge_{jk}$ genotype-by-environment interaction; $z_t$ breeding regions; $gz_{jt}$ genotype-by-region interaction.

To graphically evaluate how the data generated by the model approaches the true generative process of the observed data, we generated samples (*y*_*gen*_) from the fitted models by ancestral sampling from the conditional joint distribution and plotted against the observed data. We also accessed how the statistics maximum, minimum, median, mean and standard deviation of the data generated by the model approaches the ones of the observed data (Gelman et al. 2013, Chapter 7). For the case of maximum, this test would be: *p*_*max*_ = *Pr*(*T* (*y*_*gen*_, θ) ≥ *T* (*y*, θ)|*y*), with the same procedure using the ratio of indicators as before. In this case, the closer the Bayesian p-values are to 0.5, the higher the resemblance of the statistics from the generated versus observed data.

The Watanabe-Akaike information criterion 2 (WAIC2) was used as selection criteria to assess the goodness-of-fit of the proposed models for the maize dataset, where the lowest WAIC2 value was considered (Gelman et al. 2013, Chapter 7). The selected model was further used to compute the probabilities proposed to study GEI.

## Results

We fitted four Bayesian models for the maize dataset accounting for either homogeneous or heterogeneous residual standard deviations, as well as control for breeding region variability. All models had the 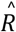 statistic close to one (see material and methods section), which indicates a convergence of the evaluated models in the parameter space. The test statistics of the observed against generated data showed all Bayesian p-values between 0.27 and 0.86 (Table 5). These results indicate that the statistics of the data generated by the model did not diverge substantially from the observed data. In addition, model M4 had the best-fit model according to the WAIC2 information criteria (Table 5). This model controlled for both heterogeneous standard deviations for environments and breeding regions. The M4 was further indicated as the best candidate model to describe the true generative process by the similarity between the densities of the observed and generated data by the model (Figure 1A).

**Table 5.**
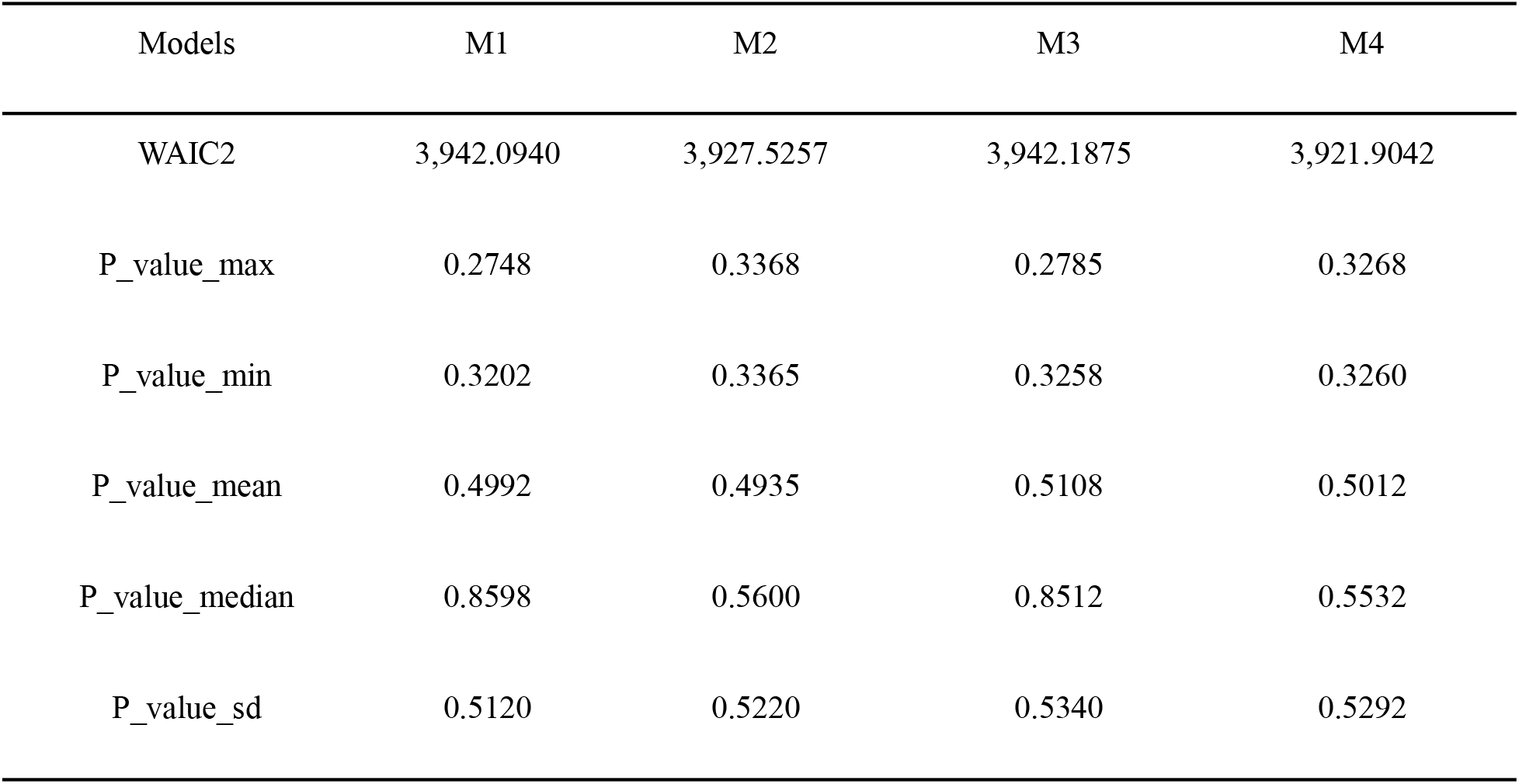
Comparative data score statistics of the Bayesian models using the maize dataset. WAIC2 shows the accuracy of the fitted models, with lower WAIC2 scores indicating better predictions.

| Models | M1 | M2 | M3 | M4 |
| --- | --- | --- | --- | --- |
| WAIC2 | 3,942.0940 | 3,927.5257 | 3,942.1875 | 3,921.9042 |
| P_value_max | 0.2748 | 0.3368 | 0.2785 | 0.3268 |
| P_value_min | 0.3202 | 0.3365 | 0.3258 | 0.3260 |
| P_value_mean | 0.4992 | 0.4935 | 0.5108 | 0.5012 |
| P_value_median | 0.8598 | 0.5600 | 0.8512 | 0.5532 |
| P_value_sd | 0.5120 | 0.5220 | 0.5340 | 0.5292 |

**Figure 1:**
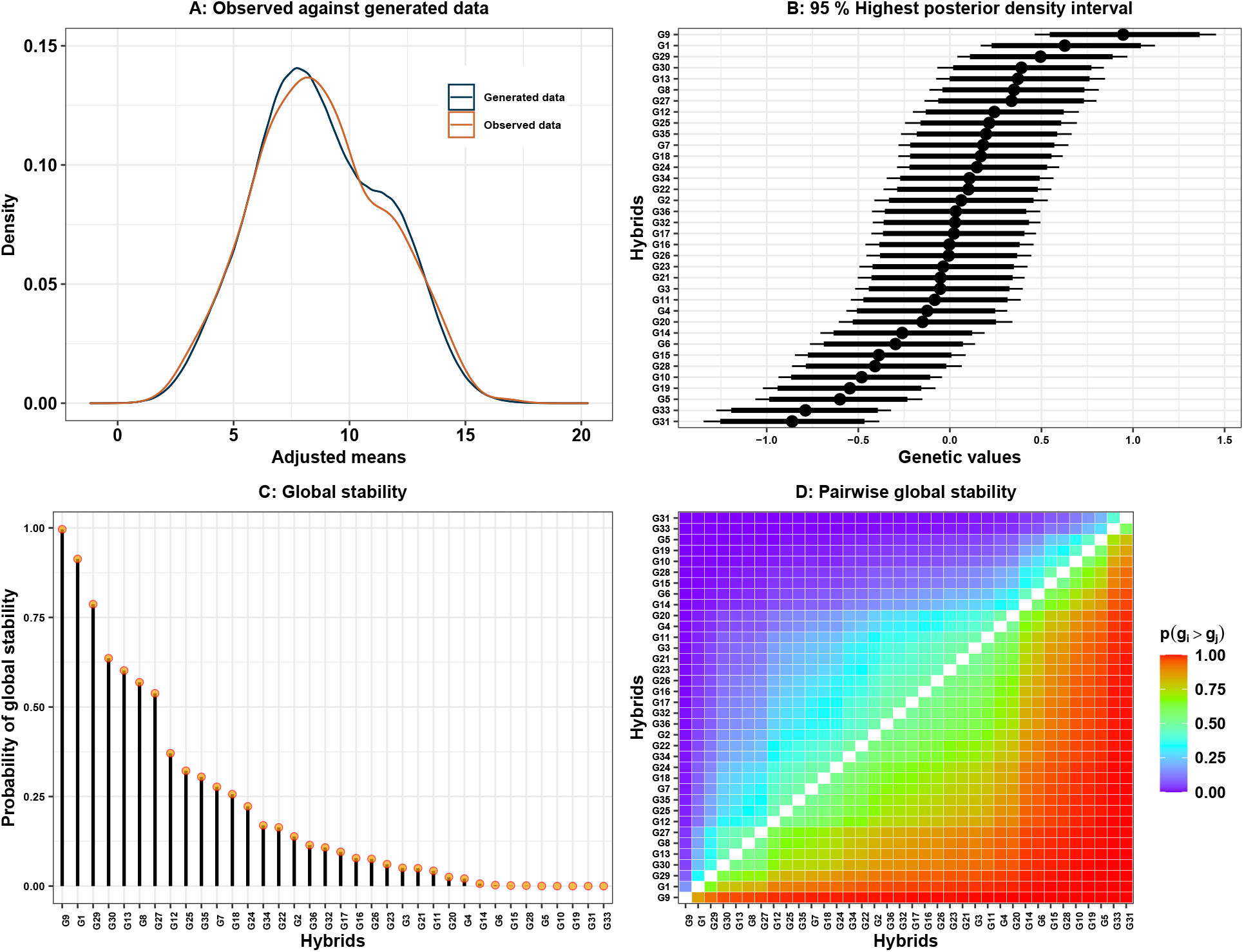
Bayesian distribution of the observed and generated data of the maize dataset described in the material and methods section (A). Caterpillar plot for hybrids posterior effects (95%) (B). The probability of a hybrid being superior to any other using the posterior effect of s Monte Carlo Sample (C). The pairwise probability among genotypes (D). All plots were based on the best selected model (M4) from Table 4.

Henceforth, considering the results of the best-fitted model (M4), the posterior distribution of the genetic values of the maize hybrids (*g*_*j*_) showed a variable overlapping pattern among their highest posterior density intervals (HDI) (Figure 1B). The hybrids G9, G1, and G29 had the highest genetic values, and the hybrids G31, G33, and G5 had the lowest genetic performance. Using the estimated genetic value, we computed the probability of performance, that is, *Pr*(*g*_*j*_ ∈ Ω | *y*) (Figure 1C). As expected, the ordering of the best performing hybrids agreed with the results from the HDI values. It also afforded a probabilistic interpretation of how likely the hybrid will be in the subset of superior hybrids based on the predefined selection intensity. All pairwise probabilities of performance are presented with a heatmap to allow comparison of the hybrid’s genetic performance (Figure 1D). In agreement with the marginal probability of performance, the hybrid G9 is also indicated as the best performing cultivar against all the others. G31 had the worst pairwise comparative performance. We also fitted seven classical models often used to study GEI with the maize dataset, named: genotypic confidence index (Annicchiarico 1992), Wricke’s ecovalence (Wricke 1965), joint regression analysis (Eberhart and Russell 1966), GGE biplot (Yan et al. 2000), AMMI method (Gauch and Zobel 1988), Shukla’s stability variance (Shukla 1972), and Lin and Binns’ superiority measure (Lin and Binns 1988). For example, in agreement with the probability of performance, G9 had the highest genetic values by the confidence index, joint regression analysis, and Lins and Binns’ superiority measure. Regarding stability, G30 was ranked among the most stable genotypes by the proposed probability stability (Figure 3B) and the well-known Shukla’s stability variance (supplementary information).

During crop improvement, essential information for selection is understanding where a potential hybrid might prevail or fail in performance. The probability of performance within environments, that is, 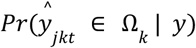, can support this breeding decision process (Figure 2A). For computing this probability using model M4, we considered the prediction 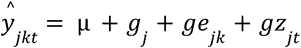, which involves genotype-zone effects. The 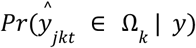 can show, for instance, that the hybrid G9, previously described with the *Pr*(*g*_*j*_ ∈ Ω | *y*) as the best-performing hybrid across environments, could fail at locations E10, E14, and E16, but is likely to succeed at the other locations (Figure 2A). Furthermore, if the location E10 is important for crop production, the hybrid G32 could also be regionally recommended in this location based on the 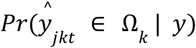.

**Figure 2:**
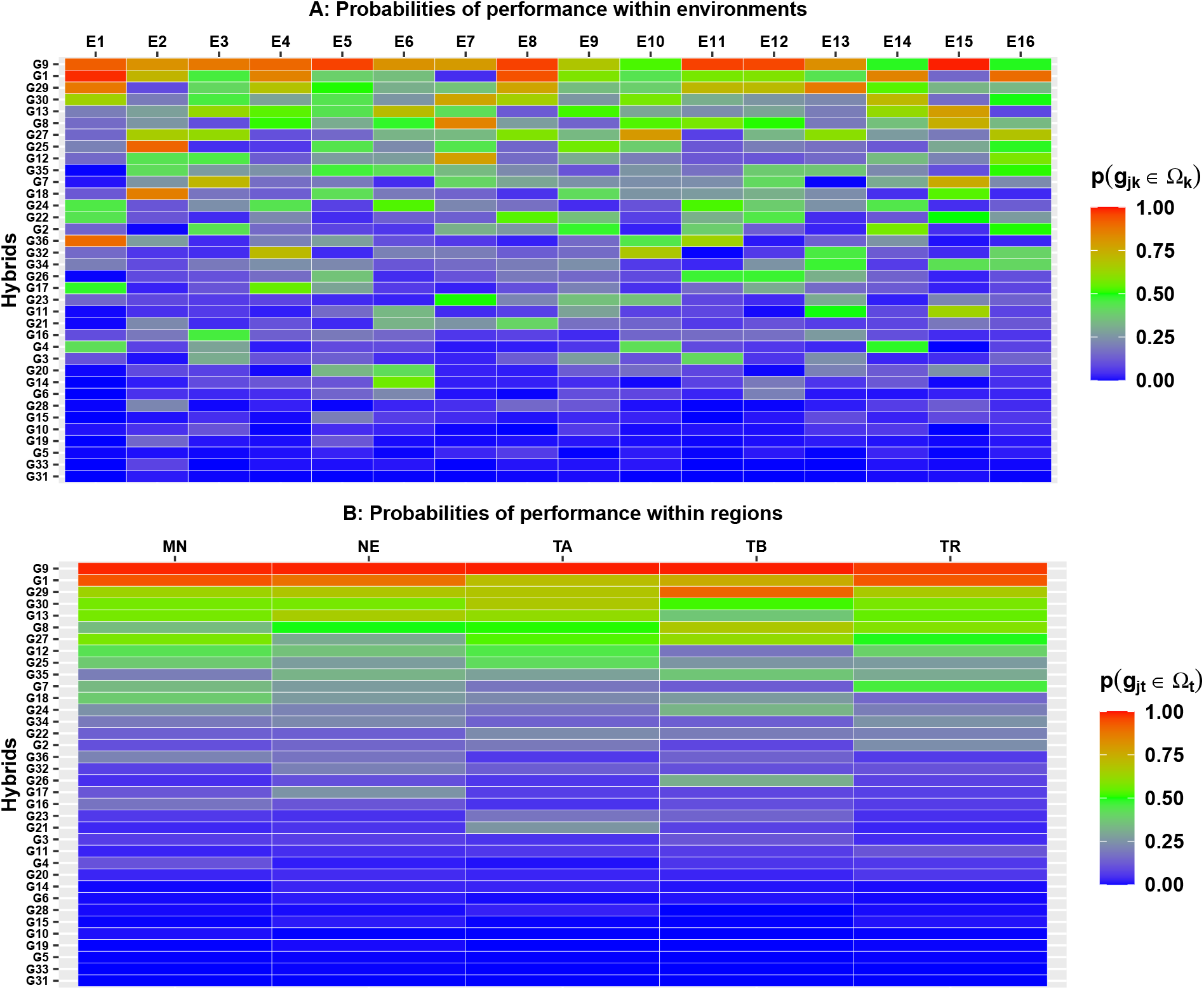
Probabilities of performance within environments (A). Probabilities of performance within breeding regions (B). Both plots were based on the best selected model (M4) from Table 4 for the maize dataset.

We can also learn more about the probability of performance within breeding regions by considering 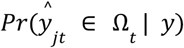, with 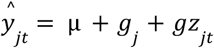 (Figure 2B). This would be important for the recommendation of cultivars to target breeding regions. For the case of the hybrid G1, we can see that 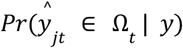 might indicate this hybrid to be cultivated in the region MN, NE, and TR with high probability, but with a lower probability for the regions TA and TB. This probability also indicates that some hybrids should not be recommended in any regions, for example, hybrids G31, G33, and G5. To identify hybrids with both high stability and yield across environments, we computed the joint probability of performance and stability, that is, 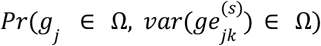 (Figure S4).

The Bayesian Eskridge’s Risk index (safety-first index), calculated with a 95% percentile with the posterior probabilities, is in agreement with our proposed joint probability of performance and stability (Figure 3), with the Spearman correlation rank equalling to 0.96. On the other hand, the rank correlation with the fixed safety-risk index (Figure S5) has a lower magnitude (Spearman correlation equal to 0.55). Based on the Bayesian Eskridge’s risk index the hybrids G9, G1, and G29 have the lower risk of recommendation across the environments (Figure 3A). Considering the computation of 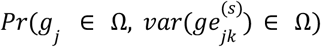 across all environments (Figure 3B), the probabilistic criteria indicated the hybrids G30, G9 and G29 had the highest stability and general performance across all environments, and the hybrids G33, G31 and G19 had lowest values of 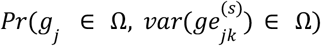.

**Figure 3:**
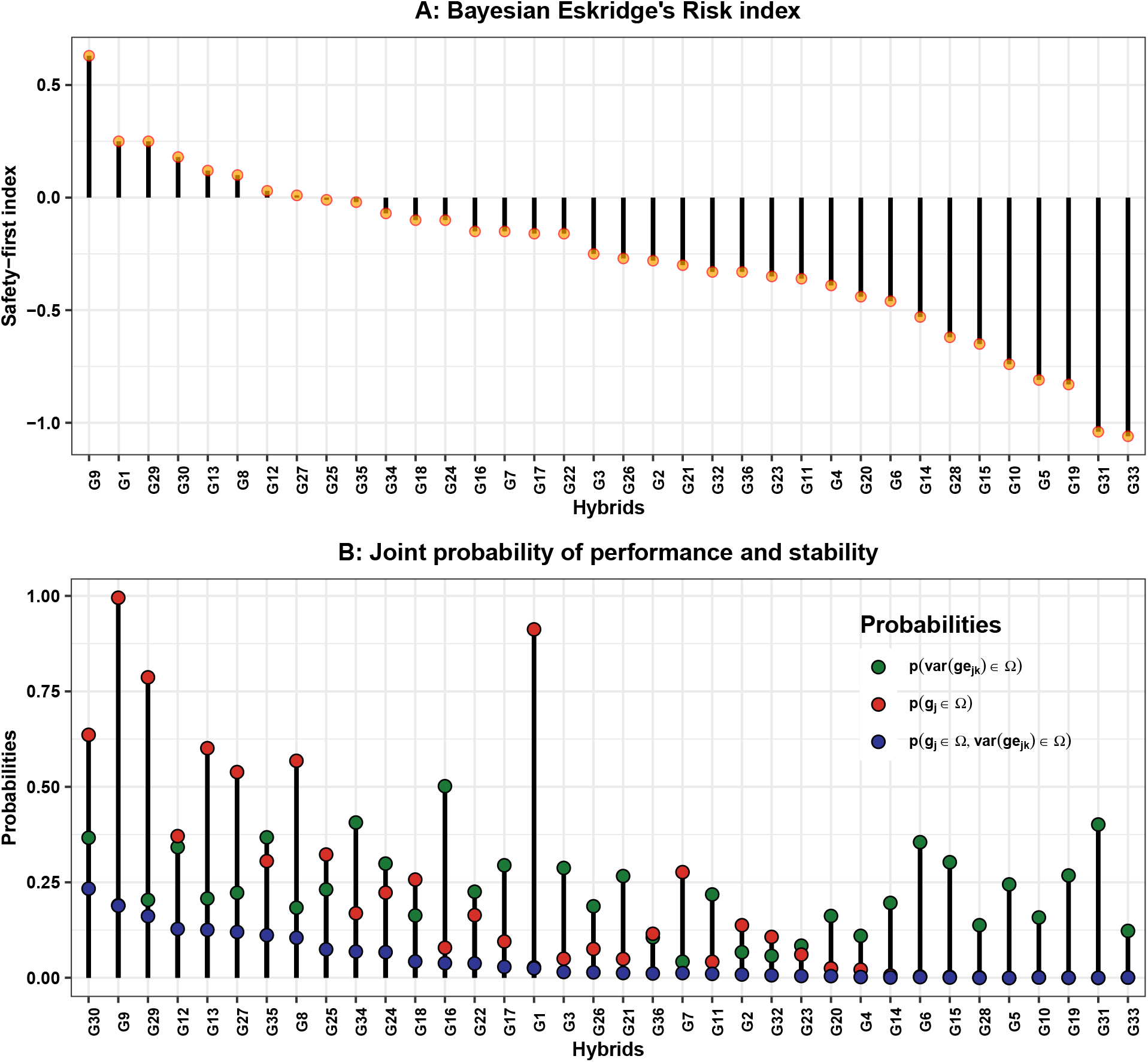
Bayesian Eskridge’s risk index calculated with a 95% percentile (A). Joint probability of performance and stability across environments (B). Both plots were based on the best-fit model M4 from Table 4 for the maize dataset.

For the wheat dataset, model M5 also presented the 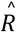 statistic closed to one with a mean value of 0.99 and a standard deviation of 0.0006. High quality of the generative process was also observed for this model (Figures S2 and S3). The probability of performance (*Pr*(*g*_*j*_ ∈ Ω | *y*)) was computed by combining the effects of breeding values across environments. Lines L463, L462, and L594 presented the highest probabilities of performance (Figure 4B). All pairwise comparisons of *Pr*(*g*_*j*_ ∈ Ω | *y*) can be assessed with the heatmap for a global comparison among lines (Figure 4C). Although line L463 presented the highest marginal probability of performance across environments (0.707), its probability of performance within the environment E1 is only 0.088. Hence, for a specific cultivar recommendation at E1, line L367 has the highest chance of success with a probability of 0.992 (Figure 4A).

**Figure 4:**
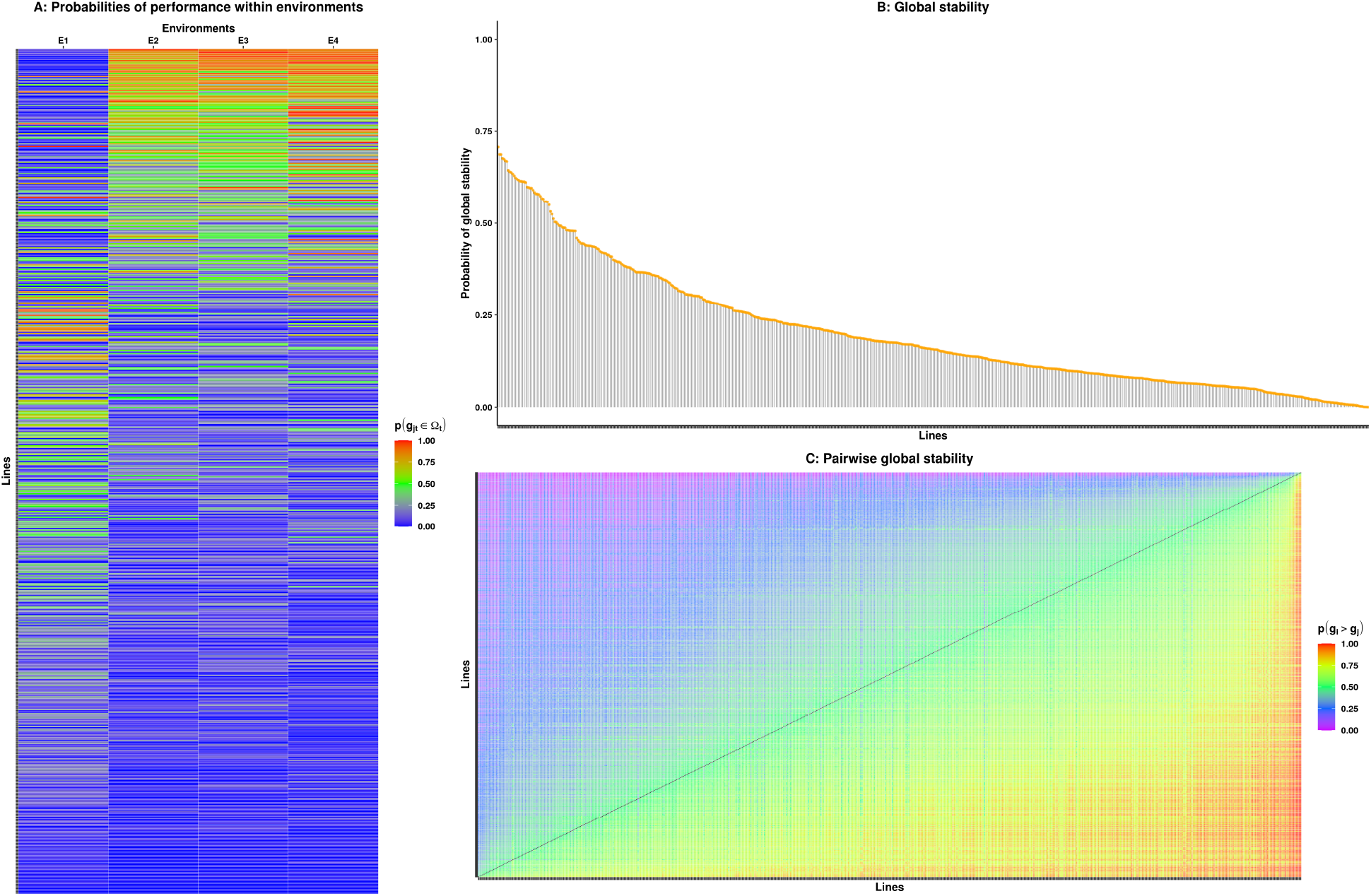
Probabilities of performance within environments (A). The probability of a line being superior to any other using the posterior effect of s Monte Carlo Sample (B). The pairwise probability among lines (C). All plots were based on the model (M5) from using the wheat dataset.

## Discussion

In this study, we proposed a Bayesian probabilistic approach to understand GEI for cultivar recommendations across a wide variety of tropical environments. For this purpose, we analyzed a maize dataset with phenotypic records of 36 single-cross hybrids evaluated in 16 target populations of environments in the tropics. The dataset contained locations covering the most crucial cropping zones in Brazil (Table S1). We also extended our proposed model to account for molecular information by using 599 lines of a wheat dataset with four environments and 1279 markers. To explore theoretical and practical concepts of applying probability methods to minimize the risk of decision-making in a breeding program, we designed non-conjugate Bayesian models for partitioning GEI. Comparison with classical GEI models showed agreement with the information provided by the probabilistic framework (supplementary information). Our approach summarizes widely used methodologies to study GEI in an easy-friendly way and provides new insights to unraveling phenotypic plasticity with probability methods.

Methodologies proposed by Mead et al. (1986), later modified by Piepho (1996), Eskridge (1990), Eskridge and Mumm (1992), and Piepho (2000) are likelihood-based approaches with additional alternatives to guide the selection by breeders. Such frequentist approaches can also be used to calculate the safety-first index (Figure S5). Eskridge and Mumm (1992) calculated the reliability of a test cultivar by taking into account normally distributed differences between 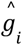 and 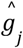 to leverage the probabilities from the cumulative function of a Normal distribution, or by considering a nonparametric model. Further probabilities/risk assessment can be estimated based on simulations assuming parameter estimates from the fitted model as true values, or in a Bayesian framework as shown in our extension for the Bayesian version of Eskridge’s Risk index (Figure 3A). Either in a frequentist or Bayesian framework, the main point of both approaches is to support breeders recommend cultivars that have little chance of having poor performance.

The proposed models take full advantage of the posterior distribution obtained with the No-U-Turn sampler algorithm to calculate the posterior probabilities. This property allows considering the uncertainty of each parameter to be estimated in a straightforward manner. We also would like to highlight that theoretically, due to the MCMC properties, as the MCMC chain sample size increases asymptotically, the probabilities approach the exact value if using the unknown cumulative posterior probability distribution from the non-conjugate Bayesian models (Goodfellow et al. 2016, Chapter 5, Gelman et al. 2013, Chapter 17).

Model selection based on WAIC2 indicated that models with heterogeneous standard deviation were substantially better than models under the homoscedastic assumption (Table 5). In addition, we could substantially improve the similarity between the densities of observed and generated data by considering the effect of the region in the model, especially for controlling different maturity zones (Figure 1A and S1). These results highlight the importance of modeling variance structures and account for different sources of variation in a MET dataset. Similar results from linear mixed models (Pastina et al. 2012; Malosetti et al. 2013) support these findings.

Model M5 leverages probabilistic concepts by exploiting genomic information by fitting a ridge regression Bayesian formulation to predict breeding values within environments. The use of molecular markers information allows predicting the performance of untested genotypes for the tested environments (Piepho et al. 2008). Thus, we can compute the probabilities of performance and stability based on prediction. These models can be further extended to deal with unbalanced historical data and covariance between environments, likewise done in a mixed model context (Piepho 1997; Smith et al. 2001; Buntaran et al. 2019; Dias et al. 2020; Krause et al. 2020) and Bayesian frameworks (Crossa et al. 2011; Josse et al. 2014; da Silva et al. 2015; dos Santos et al. 2020).

All-pairwise comparisons among the target population of genotypes are an essential task to have a better understanding of the general comparative performance between genotypes, especially in more advanced trials prior to cultivar release. Thus, the probability of performance as a function of genetic values and all-pairwise probabilities among genotypes can be used as a recommendation criterion (Figure 1C and 1D). The probabilities computed using the posterior samples can take into account the parameter uncertainty. With these measures of probabilities, the breeder can explore the probability of a given hybrid outperform any other, like the probability of a new cultivar being superior to a widespread cultivar or a reference check. As stated before, this probability has a close analogy to the Plackett-Luce model (Luce 1977; Plackett 1975). The advantage of exploring the probability of winners was reported on on-farm trials to diminish climate risks for cultivar’s recommendations (van Etten et al. 2019). Furthermore, the easy-friendly interpretation of the proposed heatmaps as a display for all-pairwise comparisons facilitates the decision-making process and allows extrapolation for other application fields.

Adaptations to new regions (mega-environments) are among the largest changes crops have gone after domestication (Wallace et al. 2018). On this basis, we computed the probability of a subset of cultivars being within the most stable based on the cultivar main effect (marginal probability) and most adaptable in a given geography (conditional probability). Most importantly, we also considered the joint distribution (product of the marginal and conditional), which capitalizes both stability and adaptation. We further estimated the probability of performance within regions to identify the genotype’s phenotypic plasticity for specific mega-environment recommendations (Figure 2B). Nonetheless, the probability of performance has the limitation of not informing about the adaptation of low-performing genotypes. Because of this potential limitation of finding stable cultivars, the stability-variance (Shukla 1972) given the intensity of selection was applied as a measure of stability 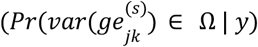. As expected, the probability of stability-variance highlighted the impact of the GEI variance of each genotype. These results guide the breeder’s decision in selecting for global or specific adaptation.

Identifying the target regions is a pivotal step to explore the genetic trends of adaptation. In this context, Bustos-Korts (2017) reported that environments with the same “winning” genotypes could be grouped as a new mega environment. Because our model explores the probability of winning given the predefined intensity of selection, the prediction of unobserved individuals in yet-to-be-seen locations might be a topic for future research.

A desired methodology that can deal with selecting for high-yielding and stable genotypes across a target population of environments has been an open question for many decades. Following this goal, a rank-sum method was proposed by Kang (1988). The method assigned ranks for stability-variance (Shukla, 1972), lowest rank receiving 1, and the highest grain yield also receiving the rank 1. Thus, the genotype with the lowest sum of the two ranks can be considered stable and high-performance. Although the rank-sum is informative, it does not provide statistical inference to compare the ranks. To quantify the risk of selection for yield and stability simultaneously, a safety-first rule was proposed in a plant breeding context by Eskridge (1990). Also, the risk index proposed by Annicchiarico (1992) takes into account cultivar means and stability simultaneously. Our approach calculates the safety-first index in a Bayesian framework (Figure 3A) by taking into account uncertainty information of the high-yielding and stability of genotypes for global or specific adaptation. For example, the joint probability of performance and stability 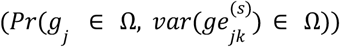 pointed out that the hybrid G9 has the highest probability of performance but does not have a high probability of stability in all breeding regions (Figure 3B). It is important to point out that the independence assumption of *g*_*j*_ and 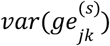 (for the maize dataset, the Pearson correlation was 0.22) could be strong and a joint probability taking into account dependence between those parameters deserves further study.

In a field trial, in general, more than one trait is evaluated and used as selection criteria. Index selection (Cerón-Rojas et al. 2008; Cerón-Rojas et al. 2016) and multi-trait models (Montesinos-López et al. 2016; dos Santos et al. 2016; Fernandes et al. 2018) have been used to get insights into how breeders should perform a selection for multiple traits. Using probabilistic methods, insights for simultaneous selection may be possible as well. For example, for drought tolerance trials, early genotypes are highly desirable. Based on a probabilistic multi-trait model, we can estimate the probability that a genotype has flowering time lower than a defined threshold. Then, we can combine this probability with the probability of performance for grain yield, resulting in a joint probabilistic score for selecting both traits. Further studies developing probabilistic scores to exploit the interaction between pleiotropic quantitative trait loci and environments could contribute to a better understanding of quantitative traits’ genetic architecture and build an index selection.

One bottleneck of selections in MET is to build reasonable criteria to define the risk of comparison among cultivars. Usually, the standard is provided by check’s performance (Eskridge and Mumm 1992), although their performance may not be equivalent across environments (Lin and Binns 1988). Given that, the latter authors proposed an index based on the distance to the maximum genotype’s response in each environment as a new standard. Our approach has the advantage that joint probabilistic models for global or specific adaptation are highly informative for decision-making based on the intensity of selection, which is more informative than environment means or distance to the maximum response. For example, in the empirical maize dataset, the next step in the breeding pipeline is selecting eight hybrids for trials of value for cultivation and use. Our model informs the uncertainty of a genotype belonging or not to the Ω for specific or global adaptation, an important piece of information in the selection process. In addition, we are addressing the GEI in favor of increasing genotype adaptation trends by doing cultivar selection and/or recommendation based on the most probable ones. Thus, we can select based on probability criteria, which is expected to increase the accuracy of cultivar selection and the potential of breeding stable crops worldwide.

It is essential to point out that the number of environments most of the time hampers the capacity of studying GEI patterns with empirical datasets. Recent advances in high-throughput phenotyping with robotic platforms and environmental covariates derived from geographic information systems are new sources of data for MET trials (Annicchiarico et al. 2006; Resende et al. 2021). Future investigation to extend our current approach with Big Data will be critical for predictive analytics to inform the performance of unobserved genotypes in yet-to-be-seen environments for the development of crops adaptable worldwide.

The main question that we addressed is the use of probability methods based on Bayesian models to improve the accuracy of cultivar recommendation for global or specific adaptation, which is crucial in any breeding program’s decision-making process. The probability of specific adaptation offers information to increase the genotype adaptation trends by allowing the identification of hybrid’s plasticity performance into specific breeding regions. Moreover, pairwise global adaptation provides a better understanding of the comparative performance between target genotypes across environments or target population of environments with an easy-friendly interpretation based on heatmaps. Finally, our findings provide a basis for unraveling genotype-by-environment interaction given a defined intensity of selection, resulting in a more informed decision-making process towards cultivar recommendation in MET.

## Author Contributions

K.O.G.D., J.P.R.S., and A.A.F.G designed the research. K.O.G.D., J.P.R.S, and M.D.K performed the statistical analysis. K.O.G.D. and J.P.R.S. wrote the first draft. A.A.F.G., M.D.K., and H.P.P critically revised drafts of the paper with editing from K.O.G.D., J.P.R.S., and M.D.K. L.J.M. and M.M.P provided the maize dataset and revised the final draft of the paper. All of the authors read and approved the paper.

## Ethics declarations

The authors declare no competing interests.

