## Supplementary material for "Leveraging probability concepts for genotype by environment recommendation": Suppemental Information (from another version of this preprint: version 1, 2021-04-22)

The supporting information is divided into the following three sections,

- **1.** Supplemental figures and tables;
- **2.** Plot and `R` code to compute Eskridge's risk index (Eskridge 1990) for the maize dataset; and
- **3.** `R` codes followed by results (figures and tables) to compute classical models often used to study GEI with the maize dataset.

### 1 Supplemental figures and tables

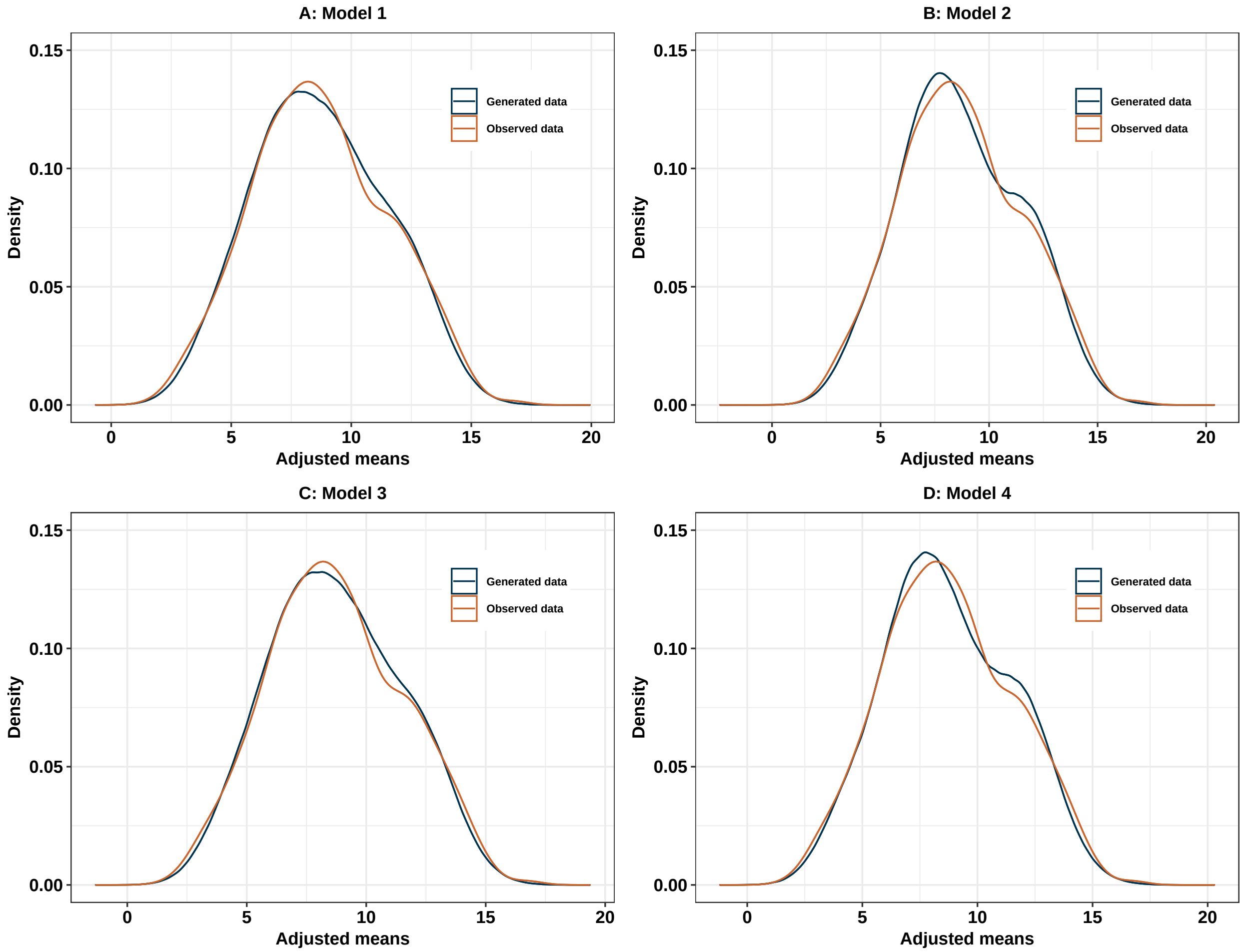


Figure S1: Bayesian distribution of the observed and generated data of the maize dataset described in the material and methods section for model M1 (A), M2 (B), M3 (C), and (M4).

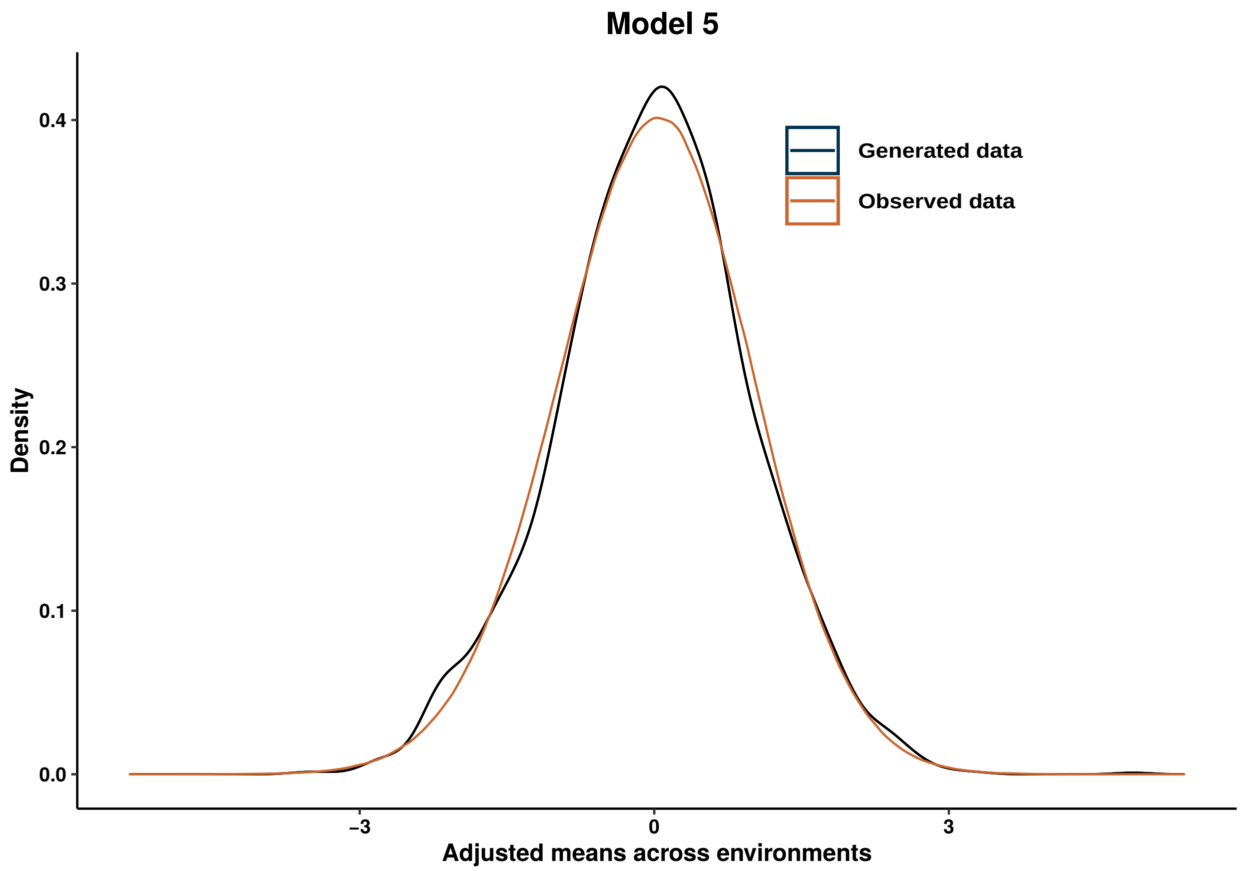


Figure S2: Bayesian distribution of the observed and generated data across environments of the wheat dataset described in the material and methods section for model M5.

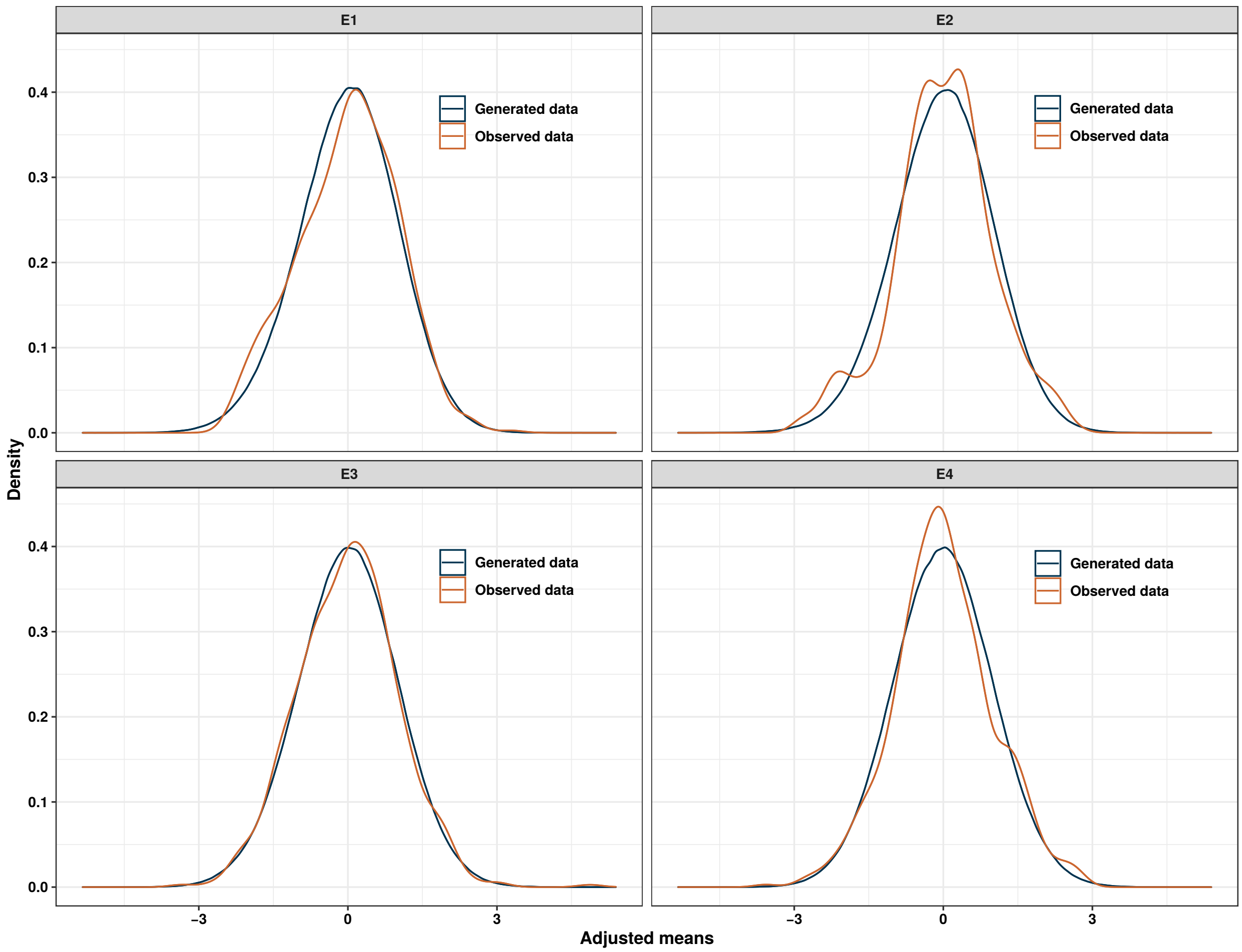


Figure S3: Bayesian distribution of the observed and generated data within environments (E1, E2, E3, and E4) of the wheat dataset described in the material and methods section for model M5.

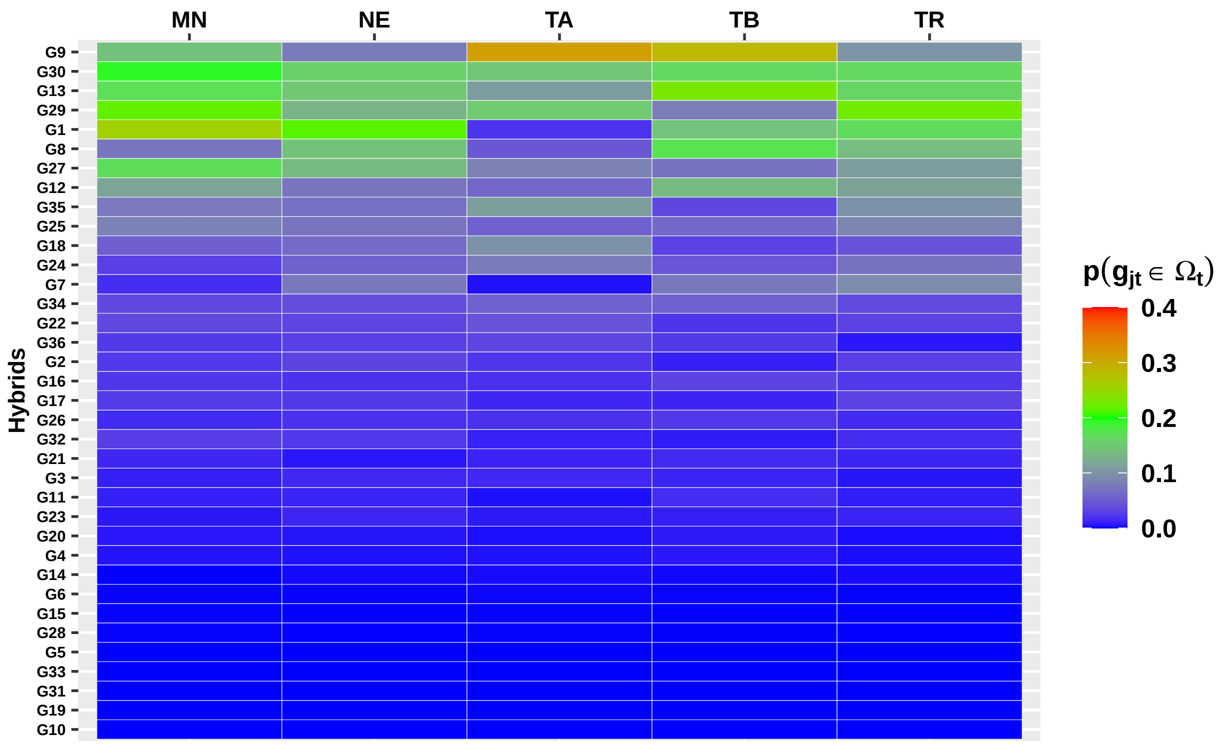


Figure S4: Joint probability of performance and stability by Regions (MN, NE, TA, TB, TR) for the maize dataset computed by model M4.

Table S1: Description of the multi-environment trials of the empirical maize dataset. CV% refers to the coefficient of variation in percentage. $H^2$ refers to generalized heritability (Cullis, Smith, and Coombes 2006).

| Region | Location | Code | Elevation | Latitude | Longitude | Mean | CV% | $H^2$ |
|---|---|---|---|---|---|---|---|---|
| MN | Mata Roma | E1 | 80 | 03º 37' S | 43º 06' W | 8.24 | 6.40 | 0.79 |
| | São Raimundo | E2 | 225 | 07º 01' S | 45º 28' W | 9.13 | 8.75 | 0.65 |
| | Teresina | E3 | 72 | 05º 05' S | 42º 48' W | 10.90 | 9.48 | 0.77 |
| NE | Nossa Senhora | E4 | 204 | 10º 29' S | 37º 11' W | 10.23 | 7.82 | 0.87 |
| | Sobral | E5 | 69 | 03º 41' S | 40º 20' W | 4.43 | 21.17 | 0.07 |
| TA | Goiânia | E6 | 749 | 16º 40' S | 49º 15' W | 6.51 | 15.83 | 0.26 |
| | Patos | E7 | 832 | 18º 34' S | 46º 31' W | 12.39 | 6.55 | 0.51 |
| | Planaltina | E8 | 944 | 15º 27' S | 47º 36' W | 13.22 | 8.06 | 0.62 |
| | Sete Lagoas | E9 | 761 | 19º 27' S | 44º 14' W | 9.37 | 13.68 | 0.44 |
| TB | Altamira | E10 | 109 | 03º 12' S | 52º 12' W | 7.06 | 14.21 | 0.31 |
| | Belterra | E11 | 152 | 02º 38' S | 54º 56' W | 9.21 | 8.50 | 0.85 |
| | Paragominas | E12 | 90 | 02º 59' S | 47º 21' W | 6.51 | 10.86 | 0.57 |
| | Rio Branco | E13 | 153 | 09º 58' S | 67º 48' W | 6.88 | 10.06 | 0.63 |
| TR | Birigui | E14 | 406 | 21º 17' S | 50º 20' W | 5.27 | 21.18 | 0.49 |
| | Campo Mourão | E15 | 585 | 24º 02' S | 52º 22' W | 11.78 | 6.37 | 0.69 |
| | Londrina | E16 | 585 | 23º 18' S | 51º 09' W | 8.26 | 13.42 | 0.54 |

### 2 Eskridge's risk index for the maize dataset

```r
# Loading the data
df <- read.csv("data.csv")

# Computing LS means within environments
install.packages("dplyr")
library(emmeans)
Envs <- levels(df$L)
results <- vector("list", length(Envs))

for (i in Envs) {
        Edat <- droplevels(subset(df, L == i))
        # Model
        mod <- aov(GY ~ H + R + B, # Obs yield ~ hybrid + replicate + block
                   data = Edat)
        temp = data.frame(lsmeans(mod, ~ H))
        results[[i]] <- data.frame(temp[,1:2], L = levels(Edat$L))
        rm(Edat, mod, temp)
}
StageI_H <- do.call(rbind, results)
StageI_H <- matrix(StageI_H$lsmean, nlevels(df$H), nlevels(df$L))

# Computing GE variance for each hybrid and marginal genotypic mean
Vii <- apply(StageI_H, 1, sd)
Yi. <- apply(StageI_H, 1, mean)

# Risk probability
Z <- qnorm(p = 0.95)
Risk_F <- round(Yi. - (Z*Vii), 2)
Risk_F <- data.frame(Risk_F, ID = levels(df$H))
Risk_F <- Risk_F[order(Risk_F$Risk_F, decreasing = TRUE),]

# Final plot
Risk_F$ID <- factor(Risk_F$ID, levels = Risk_F$ID)

x_lab <- "Hybrids"
y_lab <- "Safety-first index"

plot <- ggplot(Risk_F, aes(x = ID, y = Risk_F)) +
                geom_segment( aes(x = ID, xend = ID, y = 0, yend = Risk_F)) +
                geom_point() +
                labs(x = x_lab, y = y_lab)
```

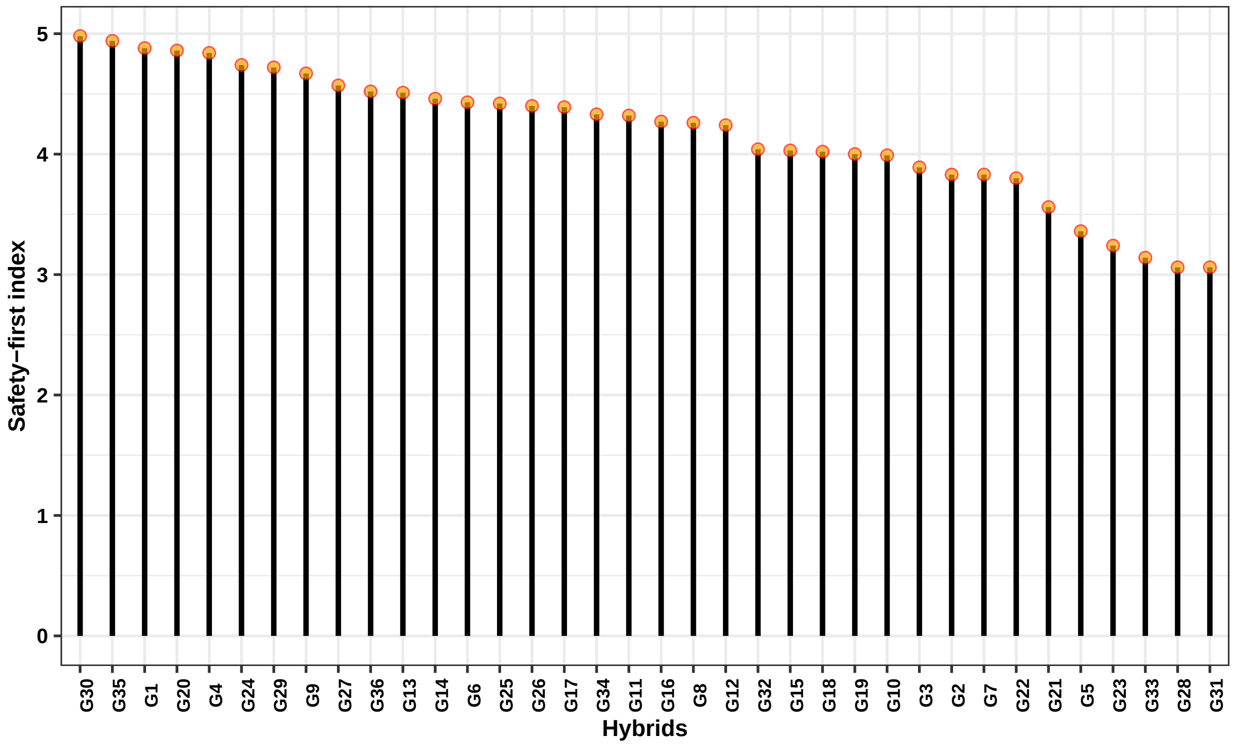


Figure S5: Eskridge's risk index calculated with a 95% percentile for the maize dataset.

### 3 Classical models often used to study GEI with the maize dataset

In order to compared the results from our proposed Bayesian model based on probability theory, we analyzed the maize dataset with the following models:

- (3.1) Genotypic confidence index (Annicchiarico 1992)
- (3.2) Wricke's ecovalence (Wricke 1965)
- (3.3) Joint regression analysis (Eberhart and Russell 1966)
- (3.4) GGE biplot method (Yan et al. 2000)
- (3.5) AMMI method (Gauch 2013)
- (3.6) Shukla's stability variance (Shukla 1972), and
- (3.7) Lin and Binns' superiority measure (Lin and Binns 1988)

The results were computed with the package `metan` (Olivoto and Lúcio 2020) in the `R` programming environment (R Core Team 2020).

#### 3.1 Installing the packages and loading the data into R memory

```
install.packages("dplyr")
library(dplyr)

install.packages("metan")
library(metan)

df <- read.csv("data.csv")
str(df)
```

```
## 'data.frame':    1152 obs. of  6 variables:
##  $ GY: num  9.65 6.72 6.67 7.69 8.99 9.53 7.28 6.42 7.24 5.72 ...
##  $ M : Factor w/ 5 levels "MN","NE","TA",..: 4 4 4 4 4 4 4 4 4 4 ...
##  $ B : Factor w/ 12 levels "1","2","3","4",..: 3 6 5 4 6 5 5 4 3 2 ...
##  $ R : Factor w/ 2 levels "1","2": 1 1 1 1 1 1 1 1 1 1 ...
##  $ H : Factor w/ 36 levels "G1","G2","G3",..: 9 20 34 18 30 27 29 26 15 19 ...
##  $ L : Factor w/ 16 levels "E1","E2","E3",..: 10 10 10 10 10 10 10 10 10 10 ...
```

Data's quality control (abnormal results, outliers identification, etc.) was already performed and is not going to be presented here. Descriptive statistics is shown in table 4 of the manuscript.

#### 3.2 Traditional visualization of genotypes in MET trials

```
ge_plot(df, gen = H, env = L, resp = GY, type = 1)
```

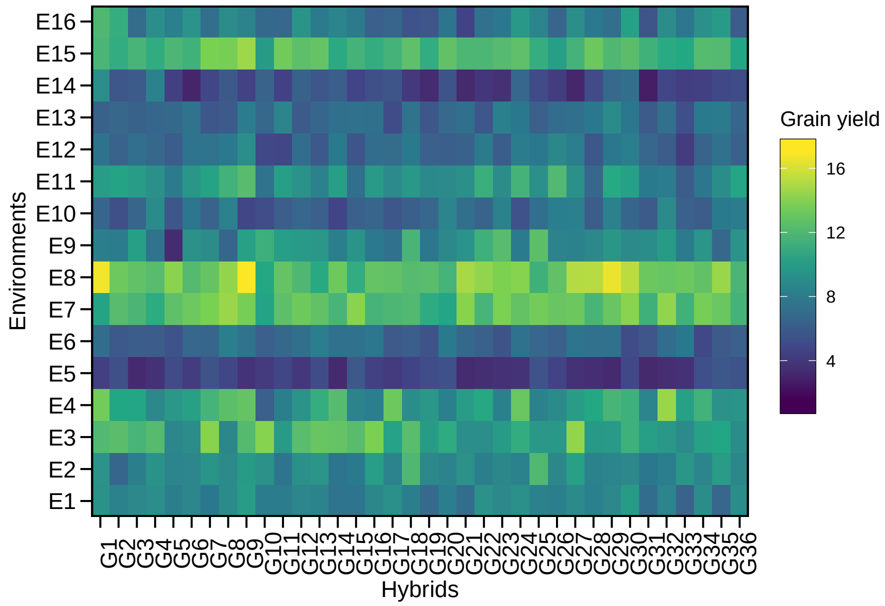


Figure S6: Heatmap of the performance of the hybrids across environments.

```
ge_plot(df, gen = H, env = L, resp = GY, type = 2)
```

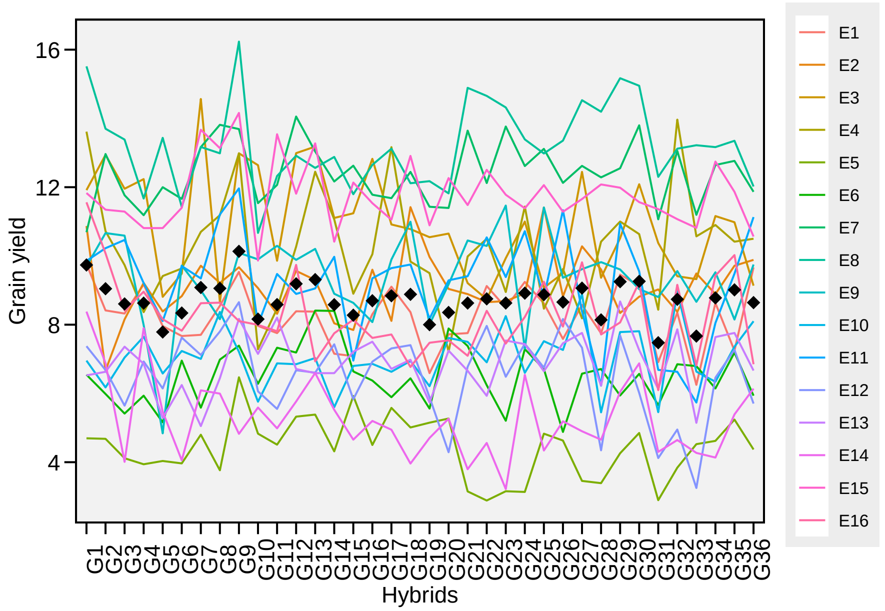


Figure S7: Reaction norm of the performance of the hybrids across environments.

#### 3.3 MET models

##### 3.3.1 Genotypic confidence index

```
ann <- Annicchiarico(df,
                     env = L,
                     gen = H,
                     rep = R,
                     resp = GY)
```

```
ann$GY$environments
```

Table S2: The mean (Y) of the environment, environmental index (index), and classification (class) as favorable and unfavorable environments (ENV).

| ENV | Y | index | class |
|---|---|---|---|
| E1 | 8.241806 | -0.4746615 | unfavorable |
| E2 | 9.132778 | 0.4163108 | favorable |
| E3 | 10.903611 | 2.1871441 | favorable |
| E4 | 10.237083 | 1.5206163 | favorable |
| E5 | 4.430417 | -4.2860503 | unfavorable |
| E6 | 6.512083 | -2.2043837 | unfavorable |
| E7 | 12.399444 | 3.6829774 | favorable |
| E8 | 13.226528 | 4.5100608 | favorable |
| E9 | 9.375972 | 0.6595052 | favorable |
| E10 | 7.062222 | -1.6542448 | unfavorable |
| E11 | 9.212500 | 0.4960330 | favorable |
| E12 | 6.518333 | -2.1981337 | unfavorable |
| E13 | 6.889306 | -1.8271615 | unfavorable |
| E14 | 5.272083 | -3.4443837 | unfavorable |
| E15 | 11.787361 | 3.0708941 | favorable |
| E16 | 8.261944 | -0.4545226 | unfavorable |

```r
ann$GY$general %>% arrange(rank)
```

Table S3: The genotypic confidence index (Wi) considering all environments, where Y is the mean yield of hybrids (GEN) across environments, its relative performance (Mean_rp) and standard deviation (Sd_rp).

| GEN | Y | Mean_rp | Sd_rp | Wi | rank |
|---|---|---|---|---|---|
| G9 | 10.133750 | 116.30498 | 13.891589 | 106.93524 | 1 |
| G1 | 9.736562 | 112.85704 | 18.422282 | 100.43140 | 2 |
| G30 | 9.261563 | 106.58340 | 9.130390 | 100.42505 | 3 |
| G12 | 9.189375 | 105.86223 | 8.215274 | 100.32111 | 4 |
| G29 | 9.250937 | 106.40029 | 10.621202 | 99.23640 | 5 |
| G13 | 9.312188 | 107.63331 | 12.888221 | 98.94034 | 6 |
| G35 | 9.012813 | 104.48673 | 9.205802 | 98.27751 | 7 |
| G8 | 9.056875 | 104.07919 | 12.692397 | 95.51829 | 8 |
| G16 | 8.696875 | 100.02364 | 7.215659 | 95.15675 | 9 |
| G2 | 9.046563 | 103.98355 | 13.895686 | 94.61105 | 10 |
| G27 | 9.063750 | 103.53054 | 13.260345 | 94.58658 | 11 |
| G7 | 9.081875 | 103.30289 | 13.413919 | 94.25534 | 12 |
| G17 | 8.842187 | 102.02845 | 11.975304 | 93.95123 | 13 |
| G34 | 8.779687 | 100.02597 | 9.114664 | 93.87822 | 14 |
| G25 | 8.873750 | 101.98165 | 12.401796 | 93.61677 | 15 |
| G18 | 8.882812 | 101.77227 | 12.641046 | 93.24602 | 16 |
| G24 | 8.917188 | 102.37851 | 13.801093 | 93.06981 | 17 |
| G11 | 8.585313 | 98.83406 | 10.566895 | 91.70680 | 18 |
| G36 | 8.642500 | 100.06012 | 12.436035 | 91.67215 | 19 |
| G26 | 8.657187 | 99.84312 | 12.140846 | 91.65425 | 20 |
| G14 | 8.581875 | 99.10956 | 12.014730 | 91.00575 | 21 |
| G3 | 8.604375 | 97.53654 | 10.553852 | 90.41808 | 22 |
| G4 | 8.627187 | 100.88326 | 16.305574 | 89.88532 | 23 |
| G6 | 8.339688 | 95.93714 | 9.563897 | 89.48639 | 24 |
| G22 | 8.751250 | 98.88815 | 14.364960 | 89.19914 | 25 |
| G21 | 8.629375 | 97.25921 | 13.249205 | 88.32276 | 26 |
| G32 | 8.738750 | 99.75522 | 17.157862 | 88.18242 | 27 |
| G20 | 8.355625 | 97.18272 | 14.445728 | 87.43922 | 28 |
| G15 | 8.276563 | 96.02864 | 13.023615 | 87.24435 | 29 |
| G10 | 8.161875 | 94.96083 | 11.910716 | 86.92717 | 30 |
| G19 | 8.000625 | 92.01892 | 9.142659 | 85.85229 | 31 |
| G23 | 8.630313 | 96.76411 | 16.346401 | 85.73863 | 32 |
| G28 | 8.139688 | 91.46935 | 13.418323 | 82.41883 | 33 |
| G5 | 7.787187 | 89.23837 | 12.579349 | 80.75373 | 34 |
| G33 | 7.668750 | 87.10158 | 15.674987 | 76.52896 | 35 |
| G31 | 7.475937 | 83.89442 | 10.975166 | 76.49178 | 36 |

##### 3.3.2 Wricke's ecovalence

```
eco <- ecovalence(df, env = L, gen = H, rep = R, resp = GY)
```

```
eco$GY %>% arrange(rank) %>% select(GEN, Ecoval, Ecov_perc, rank)
```

Table S4: Wricke's stability parameter (Ecoval) and Ecovalence in percentage value (Ecov_perc) of hybrids (GEN).

| GEN | Ecoval | Ecov_perc | rank |
|---|---|---|---|
| G19 | 13.02826 | 1.159686 | 1 |
| G34 | 13.82110 | 1.230259 | 2 |
| G35 | 14.69236 | 1.307812 | 3 |
| G16 | 14.93364 | 1.329289 | 4 |
| G30 | 15.11154 | 1.345125 | 5 |
| G31 | 15.72319 | 1.399570 | 6 |
| G12 | 15.94430 | 1.419251 | 7 |
| G6 | 17.16025 | 1.527487 | 8 |
| G3 | 19.48197 | 1.734150 | 9 |
| G29 | 22.40623 | 1.994447 | 10 |
| G14 | 25.28054 | 2.250298 | 11 |
| G15 | 25.92172 | 2.307372 | 12 |
| G11 | 26.83571 | 2.388730 | 13 |
| G26 | 27.11027 | 2.413169 | 14 |
| G24 | 27.82930 | 2.477171 | 15 |
| G17 | 29.18292 | 2.597661 | 16 |
| G21 | 29.91675 | 2.662982 | 17 |
| G18 | 30.97109 | 2.756832 | 18 |
| G13 | 32.69991 | 2.910720 | 19 |
| G27 | 33.69345 | 2.999158 | 20 |
| G22 | 33.99663 | 3.026145 | 21 |
| G28 | 34.51788 | 3.072543 | 22 |
| G9 | 35.03918 | 3.118945 | 23 |
| G36 | 35.25090 | 3.137792 | 24 |
| G8 | 36.77854 | 3.273772 | 25 |
| G25 | 37.47680 | 3.335926 | 26 |
| G20 | 38.34363 | 3.413084 | 27 |
| G2 | 38.55256 | 3.431682 | 28 |
| G5 | 39.64052 | 3.528525 | 29 |
| G23 | 40.47139 | 3.602484 | 30 |
| G33 | 42.44600 | 3.778250 | 31 |
| G4 | 42.56698 | 3.789018 | 32 |
| G7 | 43.89708 | 3.907415 | 33 |
| G10 | 43.94581 | 3.911752 | 34 |
| G1 | 62.56596 | 5.569189 | 35 |
| G32 | 66.19598 | 5.892309 | 36 |

##### 3.3.3 Joint regression analysis - Eberhart and Russell (1966)

```
jra <- ge_reg(df, env = L, gen = H, rep = R, resp = GY)
```

```
plot(jra, type = 1)
```

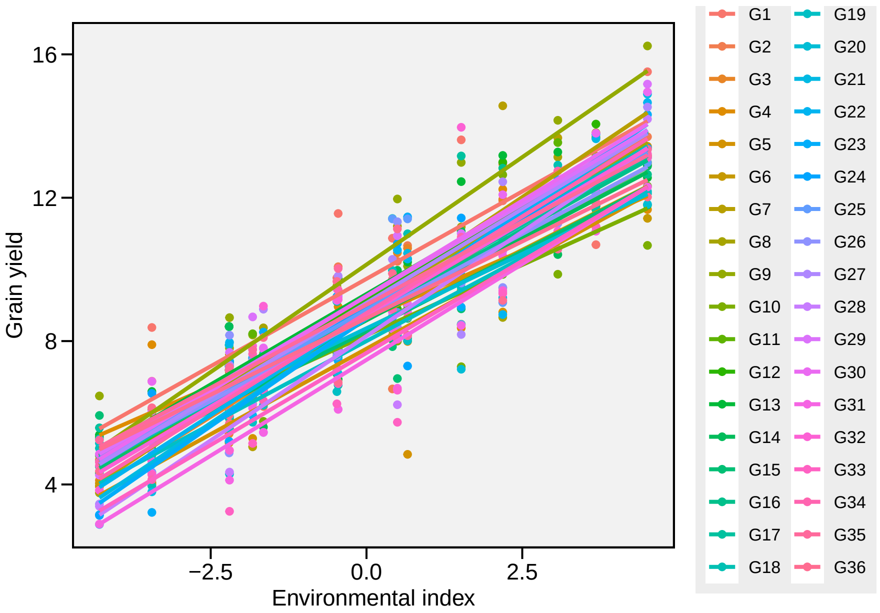


Figure S8: Plot of the regression-based stability analysis computed by the Eberhart and Russell.

```
plot(jra, type = 2)
```

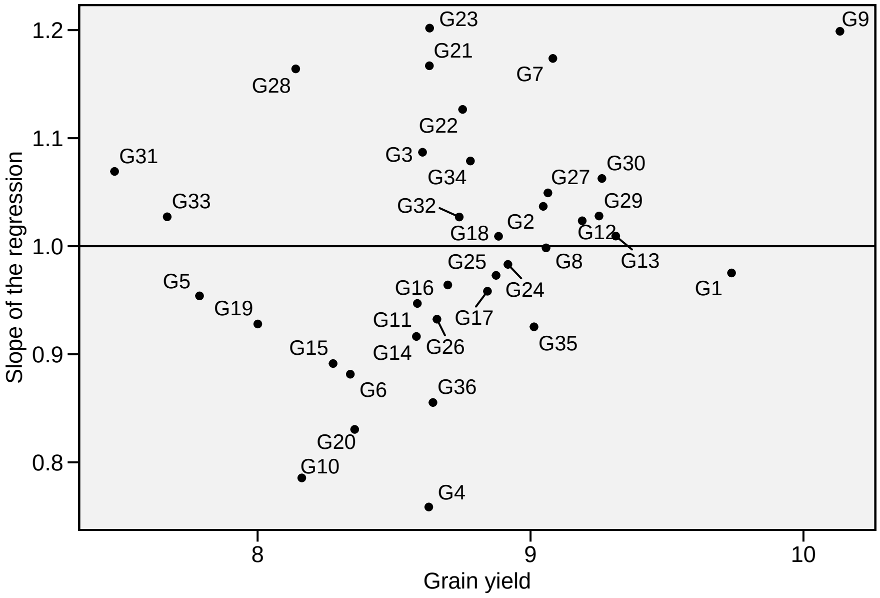


Figure S9: Slope of hybrids from the regression-based stability analysis.

##### 3.3.4 GGE biplot method

```
gge <- gge(df, env = L, gen = H, rep = R, resp = GY)
```

```
plot(gge,type = 2) ; plot(gge,type = 8) ; plot(gge,type = 6) ; plot(gge,type = 10)
```

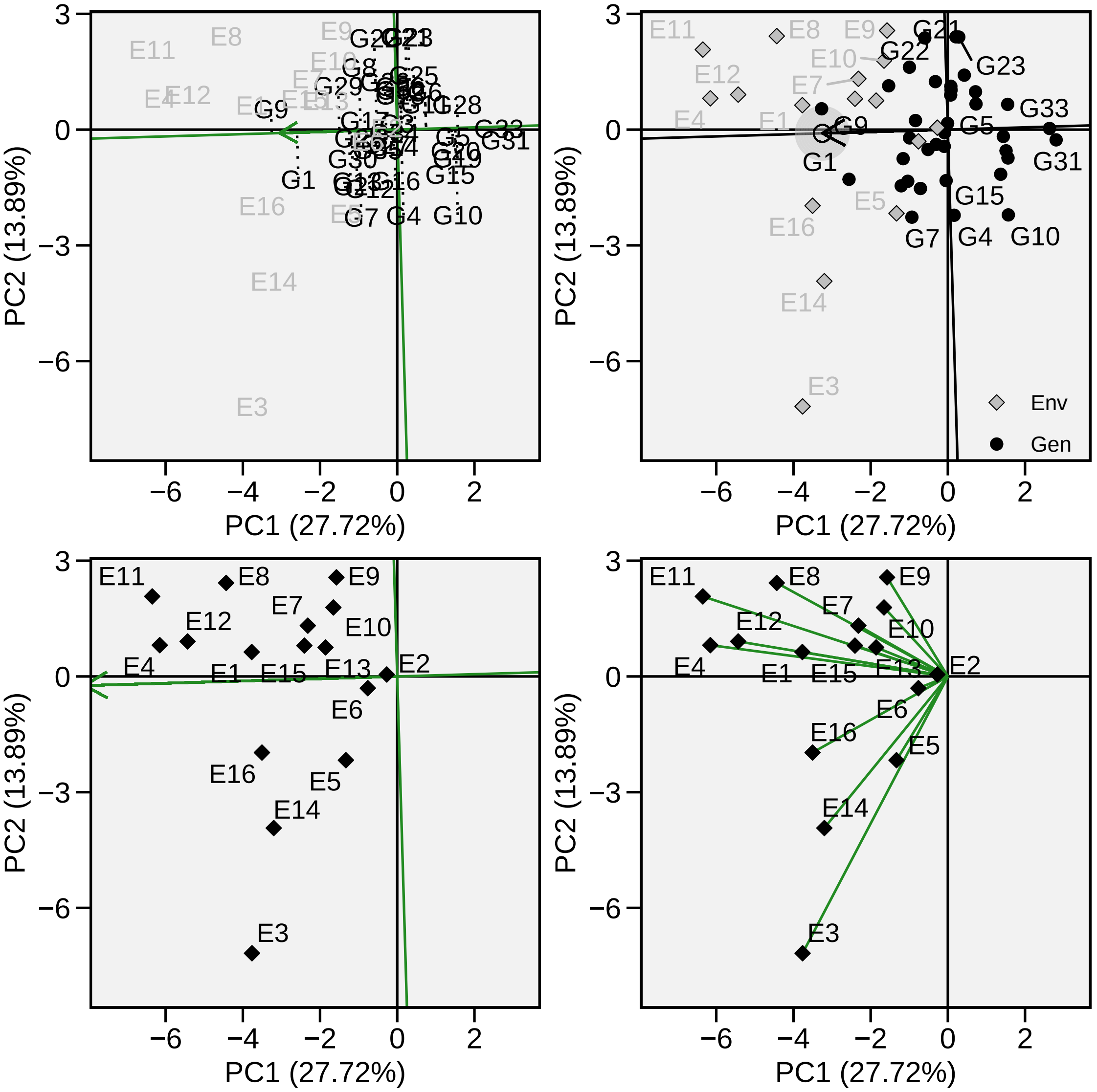


Figure S10: Plots from the GGE (G + GE) biplot model where the data was centered on environments. (a) Mean performance of genotypes versus stability; (b) Ranking of genotypes; (c) Ranking of environments; and (d) Ralationship among environments.

##### 3.3.5 AMMI model

```
AMMI <- performs_ammi(df,
                      env = L,
                      gen = H,
                      rep = R,
                      block = B,
                      resp = GY)
```

```
plot_scores(AMMI, type = 1)
plot_scores(AMMI, type = 2)
```

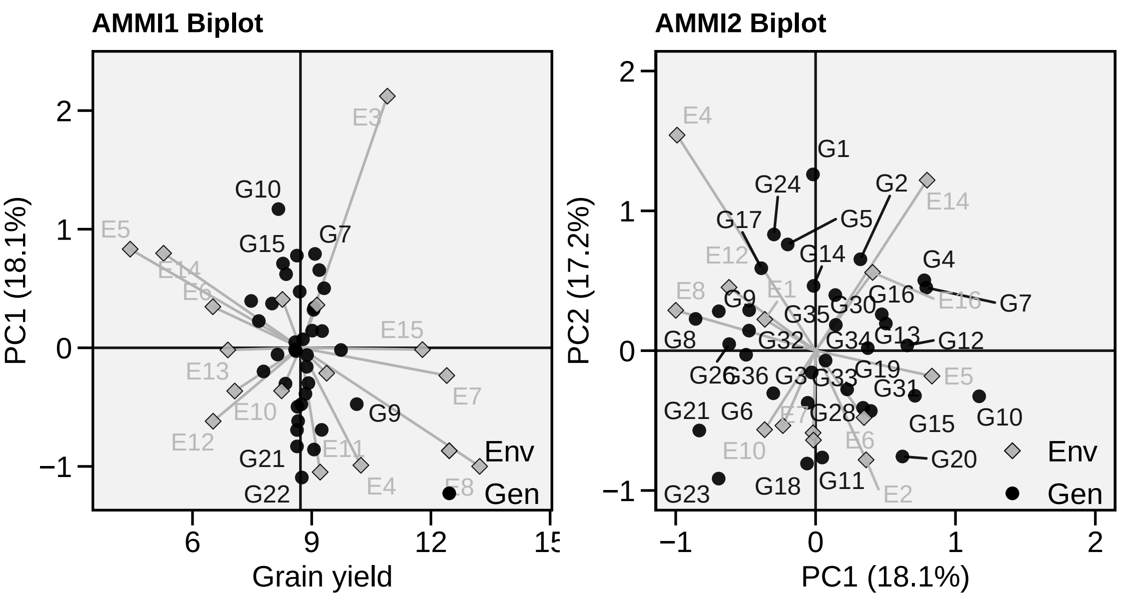


Figure S11: Plots from the AMMI model. (a) Grain yield versus first principal component (PC1); and (b) PC1 versus PC2.

##### 3.3.6 Shukla's stability variance

```
shu <- Shukla(df, env=L, gen=H, rep=R, resp=GY) ; shu$GY %>% arrange(rShukaVar)
```

Table S5: Shukla's stability variance parameter (ShuklaVar) for hybrids (GEN), where Y is the mean yield across environments, its rank (rMean), and ssiShukaVar is the simultaneous selection index (rMean + rSchukaVar).

| GEN | Y | ShuklaVar | rMean | rShukaVar | ssiShukaVar |
|---|---|---|---|---|---|
| G19 | 8.000625 | 0.4283524 | 33 | 1 | 34 |
| G34 | 8.779687 | 0.4563349 | 16 | 2 | 18 |
| G35 | 9.012813 | 0.4870853 | 11 | 3 | 14 |
| G16 | 8.696875 | 0.4956009 | 19 | 4 | 23 |
| G30 | 9.261563 | 0.5018798 | 4 | 5 | 9 |
| G31 | 7.475937 | 0.5234675 | 36 | 6 | 42 |
| G12 | 9.189375 | 0.5312713 | 6 | 7 | 13 |
| G6 | 8.339688 | 0.5741873 | 29 | 8 | 37 |
| G3 | 8.604375 | 0.6561303 | 25 | 9 | 34 |
| G29 | 9.250937 | 0.7593393 | 5 | 10 | 15 |
| G14 | 8.581875 | 0.8607855 | 27 | 11 | 38 |
| G15 | 8.276563 | 0.8834156 | 30 | 12 | 42 |
| G11 | 8.585313 | 0.9156741 | 26 | 13 | 39 |
| G26 | 8.657187 | 0.9253646 | 20 | 14 | 34 |
| G24 | 8.917188 | 0.9507418 | 12 | 15 | 27 |
| G17 | 8.842187 | 0.9985166 | 15 | 16 | 31 |
| G21 | 8.629375 | 1.0244167 | 23 | 17 | 40 |
| G18 | 8.882812 | 1.0616285 | 13 | 18 | 31 |
| G13 | 9.312188 | 1.1226458 | 3 | 19 | 22 |
| G27 | 9.063750 | 1.1577121 | 8 | 20 | 28 |
| G22 | 8.751250 | 1.1684124 | 17 | 21 | 38 |
| G28 | 8.139688 | 1.1868096 | 32 | 22 | 54 |
| G9 | 10.133750 | 1.2052082 | 1 | 23 | 24 |
| G36 | 8.642500 | 1.2126809 | 21 | 24 | 45 |
| G8 | 9.056875 | 1.2665976 | 9 | 25 | 34 |
| G25 | 8.873750 | 1.2912421 | 14 | 26 | 40 |
| G20 | 8.355625 | 1.3218358 | 28 | 27 | 55 |
| G2 | 9.046563 | 1.3292098 | 10 | 28 | 38 |
| G5 | 7.787187 | 1.3676085 | 34 | 29 | 63 |
| G23 | 8.630313 | 1.3969335 | 22 | 30 | 52 |
| G33 | 7.668750 | 1.4666256 | 35 | 31 | 66 |
| G4 | 8.627187 | 1.4708953 | 24 | 32 | 56 |
| G7 | 9.081875 | 1.5178401 | 7 | 33 | 40 |
| G10 | 8.161875 | 1.5195600 | 31 | 34 | 65 |
| G1 | 9.736562 | 2.1767417 | 2 | 35 | 37 |
| G32 | 8.738750 | 2.3048601 | 18 | 36 | 54 |

##### 3.3.7 Lin and Binns' nonparametric superiority measure

```
lab <- superiority(df, env=L, gen=H, resp = GY) ; lab$GY$index %>% arrange(R_a)
```

Table S6: Lin and Binns' superiority measure for hybrids (GEN) across all environments (Pi_a), for favorable (Pi_f) and unfavorable (Pi_u) environments (section 3.1). Y is the mean yield across environments. The indexes R* stands for the ranking for each measure (Pi_a, Pi_f, or Pi_u).

| GEN | Y | Pi_a | R_a | Pi_f | R_f | Pi_u | R_u |
|---|---|---|---|---|---|---|---|
| G9 | 10.133750 | 1.174430 | 1 | 0.5339469 | 1 | 1.814912 | 6 |
| G1 | 9.736562 | 1.539787 | 2 | 2.0189391 | 2 | 1.060634 | 1 |
| G30 | 9.261563 | 2.144354 | 3 | 2.7632953 | 5 | 1.525413 | 2 |
| G13 | 9.312188 | 2.315240 | 4 | 2.1301641 | 3 | 2.500316 | 13 |
| G12 | 9.189375 | 2.349337 | 5 | 3.0025547 | 7 | 1.696120 | 4 |
| G29 | 9.250937 | 2.450799 | 6 | 2.9575141 | 6 | 1.944084 | 7 |
| G7 | 9.081875 | 2.661984 | 7 | 2.3083938 | 4 | 3.015575 | 18 |
| G2 | 9.046563 | 2.695998 | 8 | 3.4366766 | 8 | 1.955319 | 9 |
| G35 | 9.012813 | 2.711482 | 9 | 3.7972328 | 13 | 1.625731 | 3 |
| G24 | 8.917188 | 3.060352 | 10 | 3.8833812 | 14 | 2.237323 | 11 |
| G27 | 9.063750 | 3.071120 | 11 | 4.1654250 | 17 | 1.976816 | 10 |
| G8 | 9.056875 | 3.179352 | 12 | 4.4144250 | 20 | 1.944278 | 8 |
| G17 | 8.842187 | 3.220680 | 13 | 3.6094234 | 10 | 2.831937 | 16 |
| G34 | 8.779687 | 3.285840 | 14 | 3.7434500 | 11 | 2.828230 | 15 |
| G16 | 8.696875 | 3.424381 | 15 | 4.0572516 | 15 | 2.791511 | 14 |
| G18 | 8.882812 | 3.645271 | 16 | 3.6024531 | 9 | 3.688089 | 28 |
| G3 | 8.604375 | 3.752380 | 17 | 3.7490219 | 12 | 3.755737 | 29 |
| G22 | 8.751250 | 3.774567 | 18 | 4.1233109 | 16 | 3.425823 | 26 |
| G14 | 8.581875 | 3.830009 | 19 | 4.4499047 | 21 | 3.210114 | 24 |
| G25 | 8.873750 | 3.897323 | 20 | 5.3936922 | 24 | 2.400955 | 12 |
| G26 | 8.657187 | 3.903929 | 21 | 4.7776297 | 22 | 3.030228 | 19 |
| G4 | 8.627187 | 4.046487 | 22 | 6.3509500 | 30 | 1.742023 | 5 |
| G32 | 8.738750 | 4.176084 | 23 | 5.5117891 | 25 | 2.840380 | 17 |
| G36 | 8.642500 | 4.194616 | 24 | 5.1857594 | 23 | 3.203472 | 23 |
| G21 | 8.629375 | 4.202725 | 25 | 4.2271938 | 18 | 4.178256 | 31 |
| G11 | 8.585313 | 4.315101 | 26 | 5.5734172 | 26 | 3.056784 | 20 |
| G23 | 8.630313 | 4.470440 | 27 | 4.3837172 | 19 | 4.557162 | 32 |
| G6 | 8.339688 | 4.729252 | 28 | 5.9941094 | 28 | 3.464395 | 27 |
| G15 | 8.276563 | 4.982863 | 29 | 6.7786625 | 31 | 3.187064 | 22 |
| G20 | 8.355625 | 5.118009 | 30 | 6.8195438 | 32 | 3.416475 | 25 |
| G28 | 8.139688 | 5.383977 | 31 | 5.6382031 | 27 | 5.129752 | 34 |
| G19 | 8.000625 | 5.394994 | 32 | 6.1604609 | 29 | 4.629527 | 33 |
| G10 | 8.161875 | 5.567714 | 33 | 7.9881969 | 35 | 3.147231 | 21 |
| G5 | 7.787187 | 6.512987 | 34 | 9.0693438 | 36 | 3.956630 | 30 |
| G33 | 7.668750 | 6.972638 | 35 | 7.3637672 | 33 | 6.581508 | 35 |
| G31 | 7.475937 | 7.360368 | 36 | 7.5682094 | 34 | 7.152527 | 36 |
